# The barley MLA13-AVR_A13_ heterodimer reveals principles for immunoreceptor recognition of RNase-like powdery mildew effectors

**DOI:** 10.1101/2024.07.14.603419

**Authors:** Aaron W. Lawson, Andrea Flores-Ibarra, Yu Cao, Chunpeng An, Ulla Neumann, Monika Gunkel, Isabel M. L. Saur, Jijie Chai, Elmar Behrmann, Paul Schulze-Lefert

## Abstract

Co-evolution between cereals and pathogenic grass powdery mildew fungi is exemplified by sequence diversification of an allelic series of barley resistance genes encoding Mildew Locus A (MLA) nucleotide-binding leucine-rich repeat (NLR) immunoreceptors with a N-terminal coiled-coil domain (CNLs). Each immunoreceptor recognises a matching, strain-specific powdery mildew effector encoded by an avirulence gene (*AVR_a_*_)_. We present here the cryo-EM structure of barley MLA13 in complex with its cognate effector AVR_A13_-1. The effector adopts an RNase-like fold when bound to MLA13 *in planta*, similar to crystal structures of other RNase-like AVR_A e_ffectors purified from *E. coli*. AVR_A13_-1 interacts *via* its basal loops with MLA13 C-terminal leucine rich repeats (LRRs) and the central winged helix domain (WHD). Co-expression of structure-guided MLA13 and AVR_A13_-1 substitution variants show that the receptor–effector interface plays an essential role in mediating immunity-associated plant cell death. Furthermore, by combining structural information from the MLA13–AVR_A13_-1 heterocomplex with sequence alignments of other MLA receptors, we designed a single amino acid substitution in MLA7 that enables expanded effector detection of AVR_A13_-1 and the virulent variant AVR_A13_-V2. In contrast to the pentameric conformation of previously reported effector-activated CNL resistosomes, MLA13 was purified and resolved as a stable heterodimer from an *in planta* expression system. Our study suggests that the MLA13–AVR_A13_-1 heterodimer might represent a CNL output distinct from CNL resistosomes and highlights opportunities for the development of designer gain-of-function NLRs.

## Introduction

Plant–pathogen co-evolution involves reciprocal, adaptive genetic changes in both organisms, often resulting in population-level variations in nucleotide-binding leucine-rich repeat (NLR) immune receptors of the host and virulence-promoting effectors of the pathogen^1^. NLRs often detect strain-specific pathogen effectors, so-called avirulence effectors (AVRs), inside plant cells, either by direct binding or indirectly by monitoring an effector-mediated modification of virulence targets^2^. There are two main classes of modular sensor NLRs in plants, defined by a distinct N-terminal coiled-coil domain (CC; CNLs) or a Toll-Interleukin-1 Receptor (TIR) domain, each of which plays a critical role in immune signalling after receptor activation^3, 4^. A subset of effector-activated sensor CNLs and TNLs engage additional ‘helper NLRs’ for immune signalling, some of which contain a HeLo-/RPW8-like domain or a CC at the N-terminus^5, 6^. Immune signals initiated by activated sensor CNLs, sensor TNLs and helper NLRs converge on a rapid increase in Ca^2+^ levels inside plant cells, often followed by host cell death, which is referred to as a hypersensitive response (HR)^3, 7^. In the two sensor CNLs *Arabidopsis thaliana* ZAR1 and wheat Sr35, effector-induced activation results in pentamerisation of heteromeric receptor complexes, called resistosomes, which is mainly mediated by oligomerisation of their central nucleotide-binding domains (NBDs)^8–10^. Recombinant ZAR1 and Sr35 resistosomes expressed in *Xenopus* oocytes exhibit non-selective cation channel activity, and the ZAR1 resistosome has additionally been shown to insert into planar lipid layers and display calcium-permeable cation-selective channel activity^9, 11^. Thus, currently known structures of effector-activated sensor CNLs indicate the assembly of multimeric CNL resistosomes that mediate Ca^2+^ influx in plant cells, ultimately leading to HR^3^.

In the sister cereal species barley and wheat, numerous disease resistance genes have been identified that encode CNLs conferring strain-specific immunity against the pathogenic grass powdery mildew fungi *Blumeria hordei* (*Bh*) or *Blumeria tritici* (*Bt*). Co-evolution with these Ascomycete pathogens has resulted in allelic resistance specificities at some of these loci in host populations, with each resistance allele conferring immunity only to powdery mildew isolates expressing a cognate isolate-specific AVR effector^12–16^. The *Bh* avirulence effectors AVR_A1,_ AVR_A6,_ AVR_A7,_ AVR_A9,_ AVR_A10,_ AVR_A13,_ and AVR_A22 h_ave been characterised and are recognized by the matching MLA receptors, MLA1, MLA6, MLA7, MLA9, MLA10, MLA13 and MLA22, respectively^17–19^. Although these AVR_As_ are unrelated at the sequence level, with the exception of allelic AVR_A10 a_nd AVR_A22,_ structural predictions and the crystal structure of a *Bh* effector with unknown avirulence activity (CSEP0064) suggested that they share a common RNase-like scaffold with a greatly expanded and sequence-diversified effector family in the genomes of grass powdery mildew fungi, termed RNase-like associated with haustoria (RALPH) effectors^19–22^. The crystal structures of *Bh* AVR_A6,_ AVR_A7_-1, AVR_A10 a_nd AVR_A22 v_alidated this hypothesis and revealed unexpected structural polymorphisms between them that are linked to a differentiation of RALPH effector subfamilies in powdery mildew genomes^23^. The crystal structure of the RALPH effector AvrPm2a from *Bt*, detected by wheat CNL Pm2a, was also determined and belongs to a RALPH subfamily with 34 members, which includes *Bh* AVR_A13,_ *Bh* CSEP0064 and *Bt* E-5843^16, 23^. For both barley MLA and wheat Pm2a, co-expression of matching receptor–avirulence pairs is necessary and sufficient to induce cell death in heterologous *Nicotiana benthamiana*^16–19^. Similar to several other sensor CNLs, including ZAR1 and Sr35, mutations in MLA’s MHD motif of the central NBD result in constitutive receptor signalling and effector-independent cell death (e.g., autoactive MLA10^D502V^ and MLA13^D502V^)^24–26^. While yeast two-hybrid experiments and split-luciferase complementation assays indicate direct receptor–effector interactions for several matching MLA–AVR_A p_airs, similar assays suggest that wheat Pm2a indirectly detects AvrPm2 through interaction with the wheat zinc finger protein *Ta*ZF^18, 19, 27^. The LRR of Pm2a mediates association with *Ta*ZF and recruits the receptor and AvrPm2a from the cytosol to the nucleus. However, the structural basis for how the MLA and Pm2 CNLs either directly or indirectly recognize RALPH effectors is lacking.

In this study, we used transient heterologous co-expression of barley MLA13 with its matching effector AVR_A13_-1 in *N. benthamiana* leaves and affinity purification of heteromeric receptor complexes to confirm that the effector binds directly to the receptor. In contrast to the pentameric wheat Sr35 resistosome bound to AvrSr35 of *Puccinia graminis* f sp *tritici* (*Pgt*), we find that the MLA13–AVR_A13_-1 heterocomplex is purified as a stable heterodimer and resolved using cryo-EM at a global resolution of 3.8 Å. Structural insights into the receptor–effector interface then served as a basis for structure-guided mutagenesis experiments. We co-expressed wild-type or mutant MLA13 and AVR_A13_-1 in barley leaf protoplasts and heterologous *N. benthamiana* leaves to test the relevance of effector–receptor interactions revealed by the cryo-EM structure and their roles in immunity-associated cell death *in planta*. Combining structural data with an in-depth sequence alignment between MLA receptors led to identification of a single amino acid substitution in the MLA7 LRR that allows expanded RALPH effector detection. We suggest that the stable heterodimeric MLA13–AVR_A13_-1 complex may represent an intermediate receptor– effector complex, and the equilibrium between this complex and pentameric CNL resistosomes might be differentially regulated among different sensor CNLs.

## Results

### The *in planta*-expressed MLA13-AVR_A13_-1 heterocomplex is resolved as a heterodimer

We co-expressed N-terminal GST-tagged MLA13 with C-terminal twin-Strep-tagged AVR_A13_-1 in leaves of *N. benthamiana via Agrobacterium*-mediated transformation to facilitate the formation of potential receptor–effector heterocomplexes *in planta*, followed by affinity purification for structural studies. We observed that the substitutions MLA13^K98E/K100E^, located in the CC domain, abrogate effector-triggered receptor-mediated cell death but not when MLA13^K98E/K100E^ was combined with the autoactive substitution D502V (MLA13^K98E/K100E/D502V^); Extended Data Fig. 3). Autoactivity of MLA13^K98E/K100E/D502V^ indicates that the MLA13^K98E/K100E^ substitutions do not generally disrupt receptor-mediated signalling. The MLA13^K98E/K100E^ variant allowed us to express and purify these proteins while avoiding any effect of *in planta* cell death on receptor accumulation. Analogous substitutions were introduced in the helper CNL *At*NRG1.1 which impair its cell death activity and reduces association with the plasma membrane whilst retaining oligomerisation capability^28^.

Affinity purification *via* the twin-Strep-tag on AVR_A13_-1 resulted in the enrichment of both AVR_A13_-1 and MLA13 as demonstrated by SDS-PAGE analysis (Extended Data Fig. 1). A subsequent affinity purification *via* the GST tag on MLA13 resulted in the enrichment of MLA13 with concurrent co-purification of AVR_A13_-1 (Extended Data Fig. 1), indicating that MLA13 and AVR_A13_-1 formed a heterocomplex. Further analysis of the sample by size exclusion chromatography (SEC) revealed that the heterocomplex elutes at a volume implying a molecule significantly smaller than a hypothetical multimeric MLA13 resistosome (Fig. 1a). In line with the SEC results, negative stain transmission electron microscopy (TEM) analysis revealed homogeneous particles with a diameter of approximately 10 nm, suggesting a 1:1 heterodimer of MLA13–AVR_A13_-1 rather than multimeric resistosome assemblies (Fig. 1b). Notably, star-shaped particles characteristic of pentameric resistosome assemblies such as Sr35 were completely absent (Fig. 1b).

**Fig. 1.**
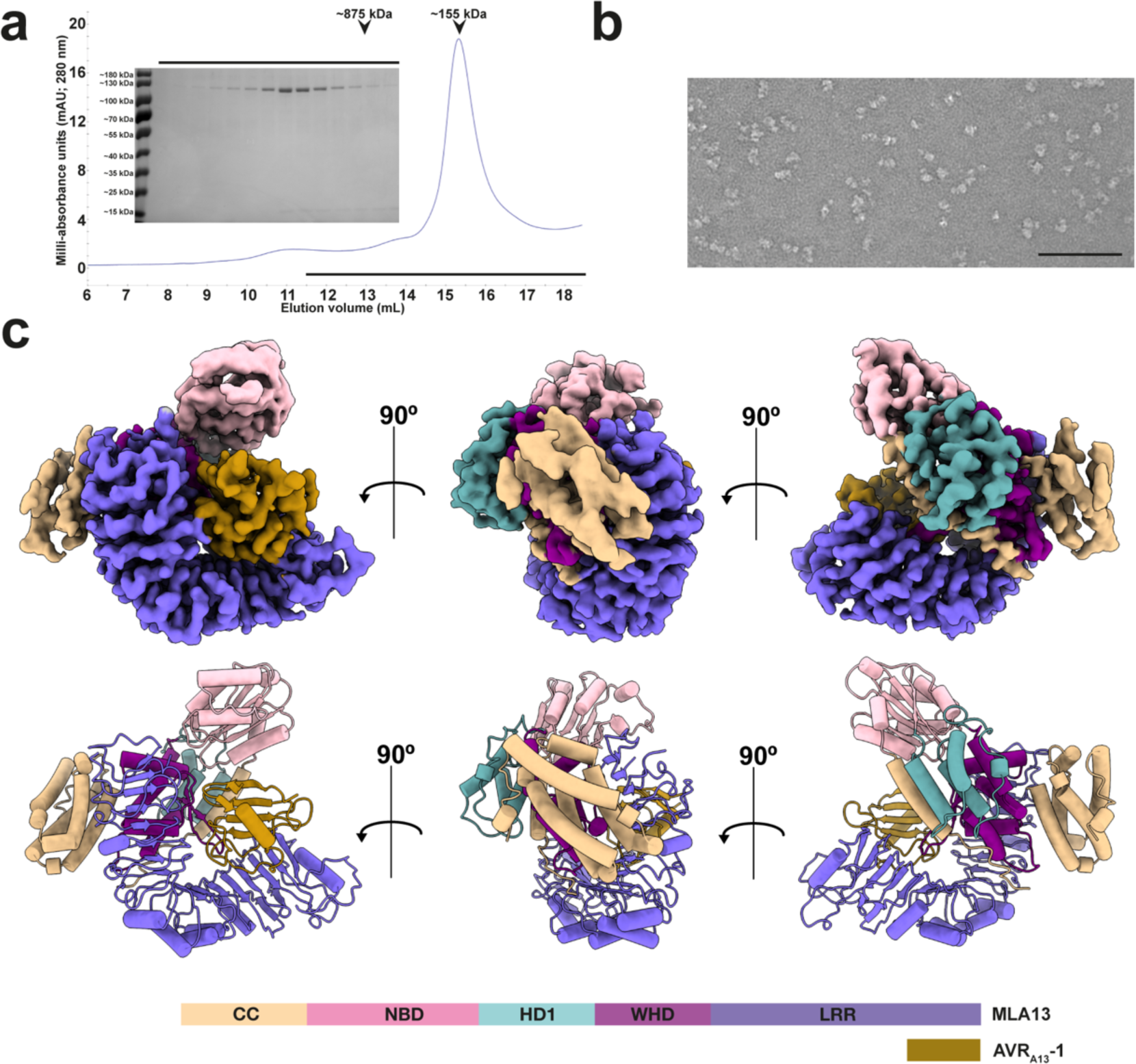
The MLA13-AVR_A13_-1 complex is purified and resolved as a heterodimer. **a,** SEC profile of the N-terminally, GST-tagged MLA13 in complex with C-terminally, twin Strep-HA-tagged AVR_A13_-1 sample purified by a two-step affinity purification as described in the Methods (Extended Data Fig.1). Inset SDS PAGE gel represents fractions eluted along the black line. The high-molecular weight marker (∼875 kDa) was determined by running the Sr35 resistosome under the same conditions. **b,** Representative negative staining image of the peak elution volume diluted five-fold. Scale bar represents 100 nm **c,** Three orientations of the MLA13-AVR_A13_-1 density map (above), atomic model (middle) and domain architecture (below). Workflow of cryo-EM data processing is presented as Extended Data Fig.2.

Previously, structures of the pentameric Sr35 resistosome were determined after co-expression of wheat Sr35 with the avirulence effector AvrSr35 of the rust fungus *Pgt* in insect cell cultures and purification of a ∼875 kDa complex by SEC^9, 10^. Stable heterodimeric MLA13–AVR_A13_-1 complex formation without detectable high-order receptor–effector complexes in *N. benthamiana* prompted us to test whether co-expression of Sr35^L11E/L15E^ with AvrSr35 in *N. benthamiana*, followed by the same purification method used for the purification of the MLA13–AVR_A13_-1 heterocomplex, leads to the formation of the Sr35 resistosome *in planta*. SEC analysis of the affinity-purified Sr35 ^L11E/L15E^–AvrSr35 heterocomplex revealed an abundant high-order complex eluting with an estimated molecular weight of 875 kDa (Extended Data Fig. 4b). Further TEM characterisation of the corresponding SEC fraction confirmed a star-shaped complex that resembles the reported insect cell-derived pentameric Sr35 resistosome^9, 10^ (Extended Data Fig. 4c). This demonstrates that the formation of the Sr35 resistosome is intrinsic to the co-expression of the two proteins, despite highly divergent expression systems in insect and plant cells. Similar results were obtained when Sr50^L11E/L15E^, an *Mla* ortholog in wheat, was co-expressed with *Pgt* AvrSr50 in *N. benthamiana*, resulting in pentameric Sr50 resistosomes upon TEM analysis (Extended Data Fig. 5)^29^. The pentameric Sr50 resistosomes purified from *N. benthamiana* are similarly star-shaped to wheat Sr35 resistosomes (Extended Data Fig.5c). In further support of these findings, blue native polyacrylamide gel electrophoresis (BN-PAGE) analysis of *N. benthamiana* leaf protein extracts provided evidence for abundant Sr35^L11E/L15E^ oligomerization when co-expressed with AvrSr35, whereas MLA13 ^L11E/L15E^ receptor oligomerization was undetectable in the presence of AVR_A13_-1 (Extended Data Fig. 6). However, oligomerization was detected when autoactive MLA13 ^L11E/L15E/D502V^ was expressed in *N. benthamiana* (Extended Data Fig.6). Collectively, this suggests that the heterodimeric MLA13-AVR_A13_-1 complex might represent an intermediate effector-activated CNL complex and that the equilibrium between heterodimeric and pentameric resistosomes may be differentially regulated among sensor CNLs. Finally, we conducted additional purification experiments to avoid potential non-native conformations, for example expression of MLA13 without an N-terminal GST tag, without substitutions in the CC domain, or equivalent mutations in the CC domains used for expressing and resolving the Sr35 and Sr50 resistosomes (Extended Data Fig.7). These experiments consistently resulted in the purification of low-order MLA13 complexes that elute from SEC at a molecular weight resembling that of the MLA13-AVR_A13_-1 heterodimer (Extended Data Fig.7).

### Cryo-EM reveals the architecture of the MLA13–AVR_A13_-1 heterodimer

Three independent MLA13–AVR_A13_-1 heterocomplex samples were prepared for cryo-EM analysis. During unsupervised 2D classification only a subset of identified particles yielded classes with features reminiscent of secondary structure elements. These had structures agreeing best with a heterodimeric but not with a pentameric assembly. Further classifying this subset of particles in 3D revealed heterodimeric complexes comprising one MLA13 and one AVR_A13_-1. Reconstruction of these particles yielded a final cryo-EM density map at a global resolution of 3.8 Å. Local resolution analysis revealed that the core region of the complex, and importantly the interface between the receptor and AVR_A13_-1, is defined up to 3.0 Å resolution. More peripheral regions such as the CC, the NBD and the first and last blades of the LRR show resolutions above 5.5 Å, implying their flexibility in the purified state of the heterodimer (Extended Data Fig. 2). Apart from these three regions, the quality of our map after machine learning-assisted sharpening was of sufficient quality to build an almost complete atomic model of the MLA13–AVR_A13_-1 heterocomplex.

The overall architecture of the MLA13–AVR_A13_-1 heterodimer resembles a single effector-bound protomer of the pentameric Sr35 resistosome^9, 10^. While the resolution of the CC domain (MLA13^1–172^) does not allow for fitting individual side-chains, it clearly shows that the four amino terminal alpha helices (α1 to α4A) form a bundle reminiscent of the ligand-bound, monomeric Arabidopsis ZAR1–RKS1– PBL2^UMP^ complex (Fig. 2a)^30^. Helix α3 is in close contact with a section of the MLA13 LRR (MLA1^518–956^) that comprises a cluster of arginine residues (MLA13^R935/R936/R559/R561/R583/R612/R657/R703^). This interdomain interaction is believed to be a precursor to formation of the ‘EDVID’ motif-arginine cluster observed in the ZAR1 and Sr35 resistosomes following activation and CC rearrangement^9, 10^. The linker (MLA13^131-143^) between helix α4A and the NBD (MLA13^173-328^) lacks observable density, suggesting significant flexibility.

**Fig. 2.**
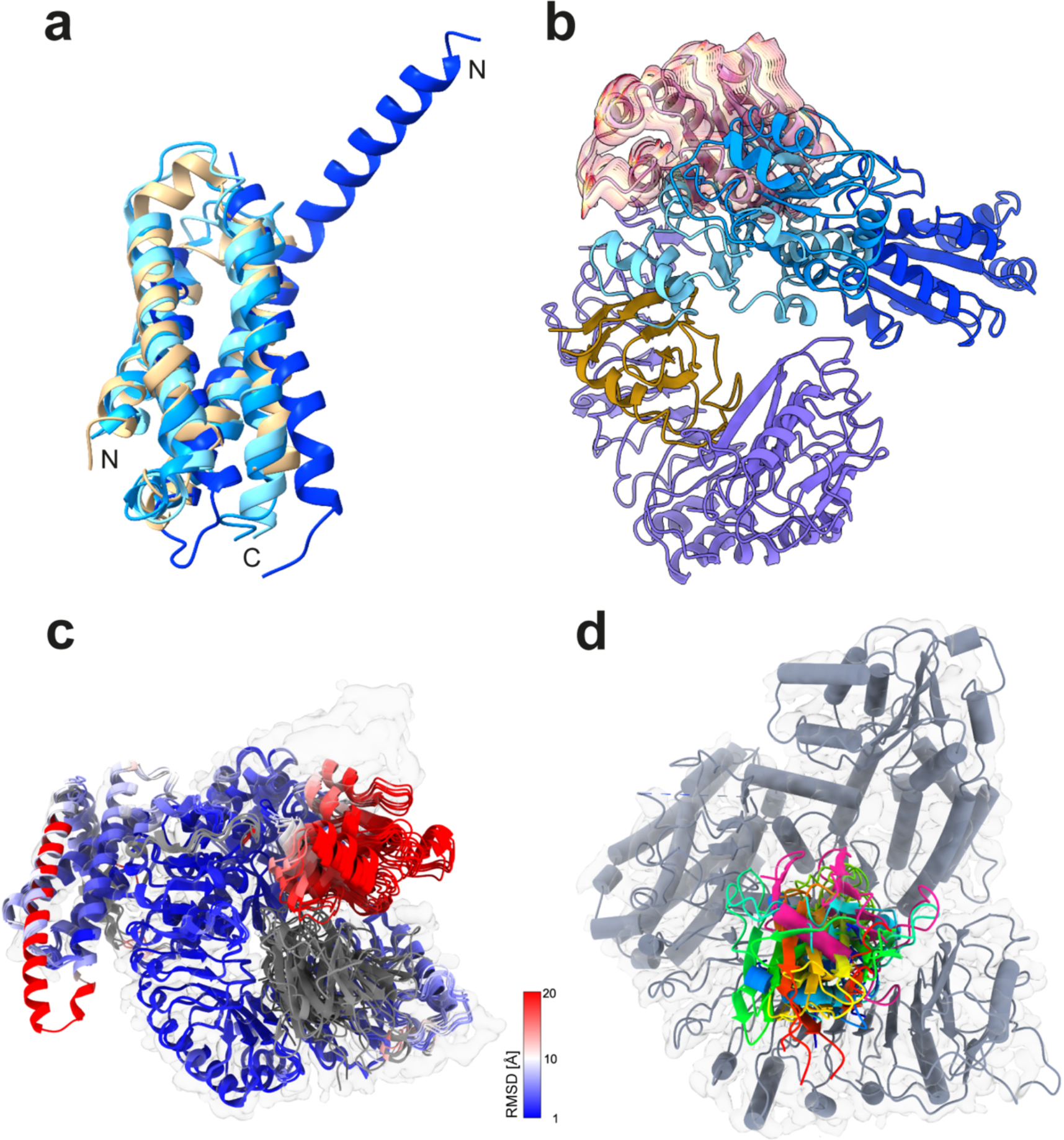
Conformational comparisons of the MLA13-AVR_A13_-1 heterodimer with ZAR1 and Alphafold predictions. **a,** Structural alignment of the CC domains of ZAR1-RKS1 (light blue; PDB: 6J5W), ZAR1-RKS1-PBL2^UMP^ (blue; PDB: 6J5V) and ZAR1 resistosome (dark blue; PDB: 6J5T) to the CC domain of the MLA13-AVR_A13_-1 heterodimer (beige). **b,** Structural alignment of ZAR1-RKS1 (light blue; PDB: 6J5W), ZAR1-RKS1-PBL2^UMP^ (blue; PDB: 6J5V) and ZAR1 resistosome (dark blue; PDB: 6J5T) to the MLA13-AVR_A13_-1 heterodimer. Only the MLA13 NBD and LRR, AVR_A13_-1 and NBDs of ZAR1 are shown. The red-yellow-red traces illustrate the major mode of conformational heterogeneity observed for the MLA13 NBD (average position shown in pink). **c,** Top five models for the MLA13-AVR_A13_- 1 complex as predicted by AlphaFold 3. All five models were aligned to the MLA13-AVR_A13_-1 experimental atomic model (grey) and predicted models are coloured by their RMSD deviation to the experimental model. For all models, the position of the NBD does not align with the experimental model. The fourth helix of the CC bundle of one predictive model is too far elongated compared to the experimental model. The experimentally observed electron density map is shown in transparent grey. **d,** AlphaFold 3 predicts five different orientations of AVR_A13_-1 (coloured rainbow) that are all incorrectly rotated compared to the experimentally observed position (pink). The MLA13-AVR_A13_-1 experimental electron density map and model are shown in transparent grey and grey, respectively.

Similar to the CC domain, the quality of cryo-EM density for the majority of the NBD does not allow for fitting individual side-chains. In addition, the canonical nucleotide binding site that is sequence-conserved with ZAR1 and Sr35 clearly lacks density for an ATP or ADP, similar to the ZAR1–RKS1–PBL2^UMP^ complex (PDB: 6J5V)^30^. This suggests that the complex might be in an intermediate state after effector binding-induced release of ADP but before ATP binding-induced oligomerisation. Overlay of the MLA13 NBD after AVR_A13_-1 binding to the receptor with the NBD of an Alpha-fold3 model of the AVR_A13_-1-bound MLA13 receptor shows conformational differences in NBD conformations between the prediction and experimental model (Fig. 2c). In addition, a motion-based deep generative model to investigate the flexibility remaining in the subpopulation of particles used for the 3D refinement implies that the NBD can sample a conformational space by rotating relative to the WHD (MLA13^410-517^) (Fig. 2b). Interestingly, a similar hinge situated between the NBD and the WHD domain is observed when comparing the MLA13 NBD position to the NBD position in ZAR1 bound or unbound to the effector^30^. Despite its flexibility, the MLA13 NBD does not, however, sample positions overlapping with the ZAR1 NBD, and the consensus position is about 75 degrees rotated compared to the ZAR1 resistosome (Fig.2b). Despite the differences observed for the NBD, the remaining domains of MLA13, namely HD1 (MLA13^329-409^), WHD, and LRR, adopt positions similar to those observed in the non-resistosome ZAR1 structures (PDBs: 6J5W and 6J5V)^30^.

### AVR_A13_-1 adopts an RNase-like fold *in planta* and interacts both with the LRR and the WHD domain of MLA13

AVR_A13_-1 adopts an RNase-like fold reminiscent of the crystal structures reported for *E. coli*-expressed AVR_A6, A_VR_A7_-1, AVR_A10 a_nd AVR_A22 o_f *Bh*, all of which share a structural core of two β-sheets and a central α-helix (Fig. 3a)^23^. The N-terminal β-sheet consists of two antiparallel strands (β1 and β2), whilst the second β-sheet consists of four antiparallel β-strands (β3 to β6). Based on structural polymorphisms between *Bh* AVR_A6, A_VR_A7_-1, AVR_A10,_ AVR_A22 a_nd *Bt* AvrPm2, AVR_A13_-1 is most similar to *Bt* AVR_Pm2 a_nd the structure of a *Bh* effector with unknown avirulence activity, CSEP0064^21, 23^. Each of the four crystallised AVR_A e_ffectors and *Bt* AvrPm2 share two conserved cysteine residues at the N and C termini, respectively, that form an intramolecular disulphide bridge connecting the N- and C-terminals. In AVR_A13_-1, however, the position of the N-terminal cysteine is occupied by a leucine, preventing intramolecular disulphide formation with the C terminal residue AVR_A13_-1^C116^ (Fig. 3a). The conserved structural core of AVR_A13_-1 and proximity of AVR_A13_-1 N- and C-terminal ends show that intramolecular disulphide bridge formation is likely dispensable for adoption of an RNase-like fold when bound to its receptor inside plant cells (Fig. 3a). This also indicates that binding to the receptor does not lead to extensive rearrangements of the RNase-like fold compared to AVR_A c_rystal structures of proteins purified from *E. coli* and unbound to their matching receptor^22^.

**Fig. 3.**
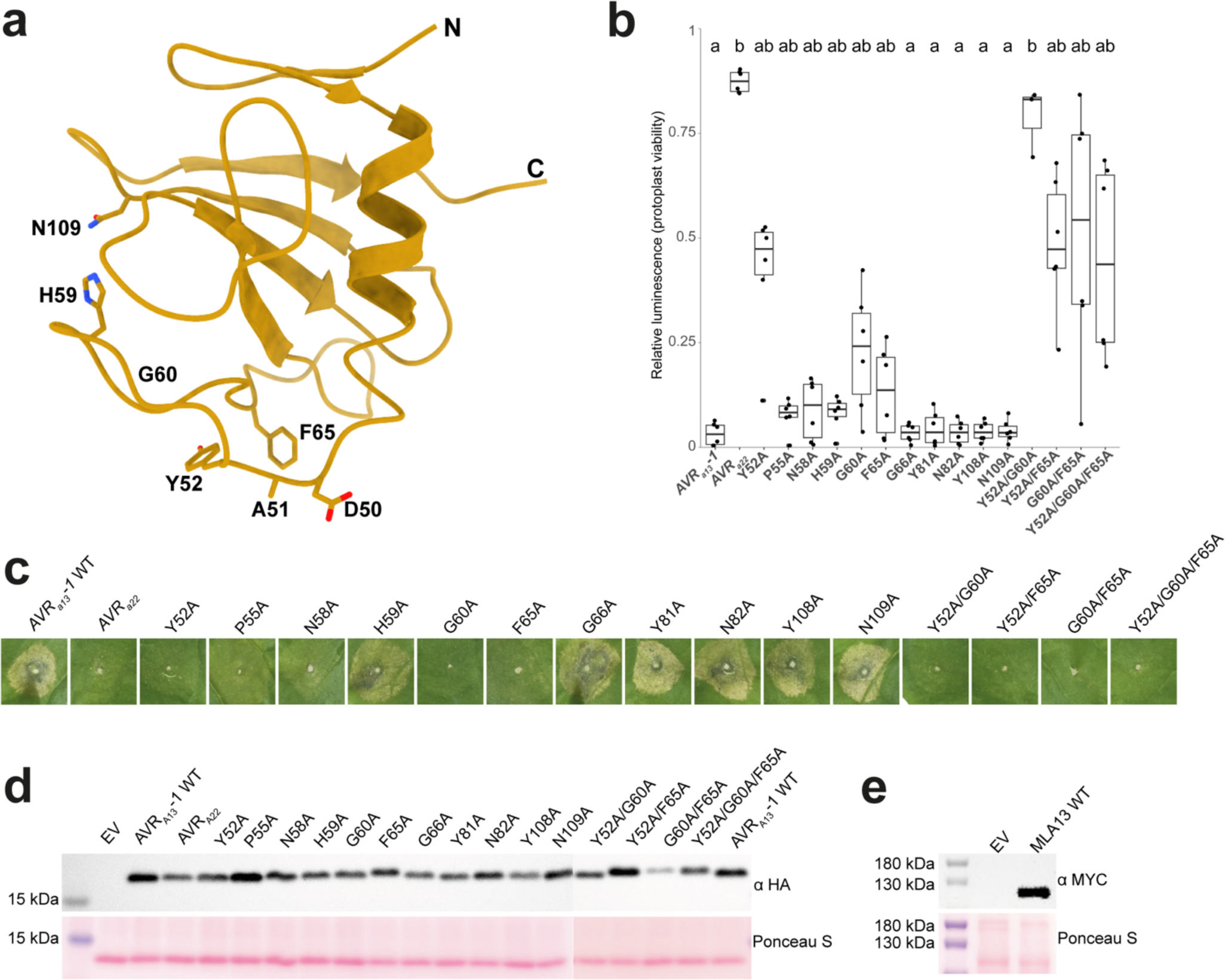
The AVR_A13_-1 basal loops are primarily responsible for interacting with the MLA13 LRR. **a,** The cryo-EM structure of AVR_A13_-1 residues (atom display) that were experimentally shown to be responsible for triggering MLA13-mediated cell death. **b,** Co-expression of MLA13 with AVR_A13_-1 substitution mutants in barley protoplasts. Luminescence is normalised to EV + MLA13 (= 1). High relative luminescence suggests low cell death response and therefore suggesting loss of AVR_A13_-1 interaction with MLA13. The six data points represent two technical replicates performed with three independently prepared protoplast samples. Treatments labelled with different letters differ significantly (*p* < 0.05) according to the Dunn’s test. **c,** *Agrobacterium*-mediated co-expression of MLA13 with AVR_A13_-1 interface substitution mutants in leaves of *N. benthamiana*. Three independent replicates were performed with two *Agrobacterium* transformations and plant batches (Supplementary Fig. 2). **d,** Western blot analysis of AVR_A13_-1 substitution mutants. **e,** Western blot analysis of MLA13.

The cryo-EM density with higher local resolution of the interface between the MLA13 LRR and AVR_A13_-1 reveals interactions of the effector with multiple receptor residues, specifically from the concave side of the LRR and the WHD (Fig. 4a). To investigate the physiological relevance of the interactions between MLA13 and AVR_A13_-1, we generated substitution variants of putative interacting residues in both the receptor and effector; we then transiently expressed these in barley protoplasts and leaves of *N. benthamiana* and tested for loss of AVR_A13_-1-triggered and MLA13-mediated cell death.

**Fig. 4.**
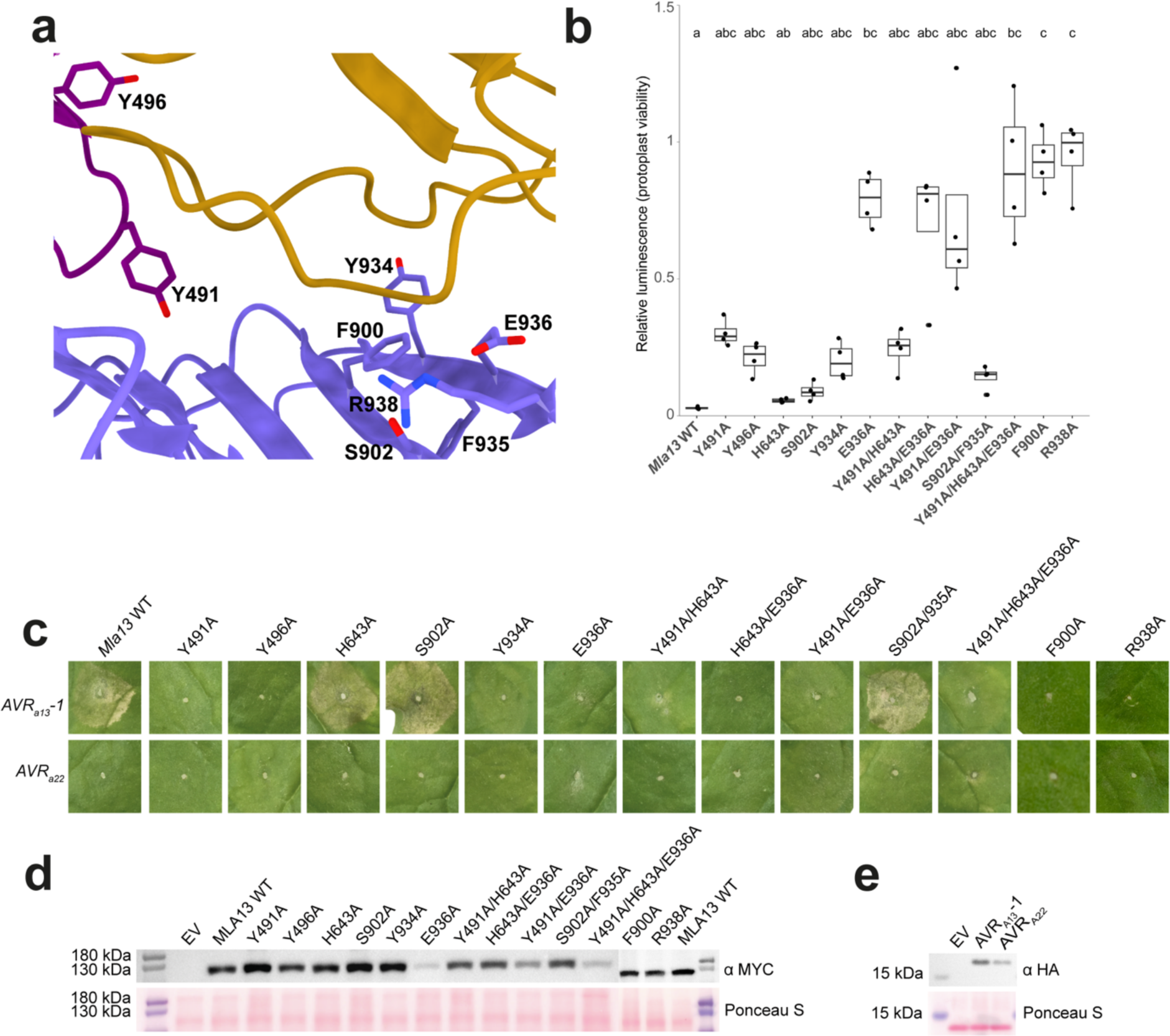
Minimal but spatially distributed recognition of AVR_A13_-1 by the MLA13 LRR and WHD. **a,** The MLA13-AVR_A13_-1 interface exhibiting MLA13 residues that were experimentally shown to contribute to AVR_A13_-1-mediated cell death. **b,** Co-expression of AVR_A13_-1 with MLA13 substitution mutants in barley protoplasts. Each MLA13 variant was normalised to its own autoactivity; luminescence is normalised to EV + MLA13 variant (= 1). High relative luminescence suggests low cell death response and therefore loss of AVR_A13_-1 interaction with MLA13. The four data points represent two technical replicates performed with two independently prepared protoplast samples. Treatments labelled with different letters differ significantly (*p* < 0.05) according to the Dunn’s test. **c,** *Agrobacterium*-mediated co-expression of AVR_A13_-1 with MLA13 substitution mutants believed to contribute to MLA13 interface and cell death response in leaves of *N. benthamiana*. Three independent replicates were performed with two *Agrobacterium* transformations and plant batches (Supplementary Fig. 3). **d,** Western blot analysis of MLA13 substitution mutants. **e,** Western blot analysis of AVR_A13_-1 and AVR_A22._

Visualisation of the MLA13–AVR_A13_-1 interface clarifies that the two basal loops of AVR_A13_-1 (AVR_A13_-1^W47-T74^) play an essential role in the interaction with MLA13 and receptor-mediated cell death. Notably, the aromatic ring from AVR_A13_- 1^Y52^ presents strong π-π stacking with MLA13^F900^ and interacts with MLA13^F934^, an observation supported by a loss in cell death activity due to the single AVR_A13_-1^Y52A^ and MLA13^F900A^ substitutions (Figs. 3b,c and 4b,c). Contributing to stabilisation of the AVR_A13_-1 basal loops and their interaction with the receptor, AVR_A13_-1^F65^ seemingly engages in a T-shaped interaction with the aromatic ring of MLA13^Y934^. Furthermore, a notable reduction of cell death was observed when stacking the two substitutions AVR_A13_-1^Y52A/G60A^, presumably generating a steric clash between the backbone of AVR_A13_-1^G60^ and MLA13^Y491^ (Fig. 3b,c and Fig. 4b,c). Reciprocally, the substitutions MLA13^Y491A^ and MLA13^Y496A^ in the WHD resulted in a reduced cell death, suggesting that the WHD plays a critical role in triggering conformational changes in MLA13 that are necessary for cell death activity (Fig. 4b,c). Additional charged π interactions between MLA13^H643^ and AVR_A13_-1^N82^ are also thought to be an important component of the receptor–effector interface. This is supported by the near-complete loss of cell death activity of the double substitution mutant MLA13^H643A/E936A^ (Fig. 4b,c). We then tested the cell death activity of individual MLA13^E936A^ and MLA13^S902A^ variants (Fig. 4b,c). While MLA13^S902A^ retained wild-type-like activity, the single receptor substitutions MLA13^F900A^ and MLA13^E936A^ resulted in a complete loss of cell death (Fig. 4b,c). Finally, we inferred that MLA13^S902^ acts to stabilise MLA13^R938^, an essential interactor of AVR_A13_-1^D50^ and AVR_A13_-1^A51^ that leads to a complete loss of cell death when introducing the single substitution MLA13^R938A^ (Fig. 4b,c).

### Expansion of MLA7 effector recognition specificity

Understanding the roles of receptor residues in the MLA13–AVR_A13_-1 interface allowed us to generate a gain-of-function (GoF) MLA receptor based on amino acid sequence alignment with known MLA resistance specificities to *Bh* (Extended Data Fig. 8)^12^. In this alignment, we observed that MLA7 is most similar to MLA13 with over 93% sequence conservation among the two LRR domains (Extended Data Fig. 8)^23^. Closer inspection of the MLA7 and MLA13 sequence alignment revealed that only one of the LRR residues contributing to the MLA13–AVR_A13_-1 interface was polymorphic between the two receptors at positions MLA7^L902^ and the corresponding MLA13^S902^ (Extended Data Fig.8). We then introduced the substitution MLA7^L902S^ to test if this MLA13-mimicking receptor could gain detection of AVR_A13_-1 while retaining the ability to detect its previously described cognate AVR_A7 e_ffectors^18^. The co-expression of MLA7 WT with AVR_A7_-2 in barley protoplasts results in a cell death response, whilst only weakly recognising AVR_A7_-1, AVR_A13_-1 and AVR_A13_-V2, a virulent variant of AVR_A13_-1 (Fig. 5a)^17, 26^. We then performed the same experiment with the MLA7^L902S^ variant: not only was cell death activity retained upon co-expression with AVR_A7_-2, but a gain of cell death activity was detected upon co-expression with AVR_A7_-1, AVR_A13_-1 and AVR_A13_-V2, a virulent variant of AVR_A13_-1 (Fig. 5a). Notably, MLA7^L902S^ does not detect AVR_A22,_ indicating that the detection GoF receptor could be limited to a subset of RALPH effectors (Fig. 5a). The same co-expression experiments were performed in leaves of *N. benthamiana* with qualitatively similar results (Fig. 5b,c,d).

**Fig. 5.**
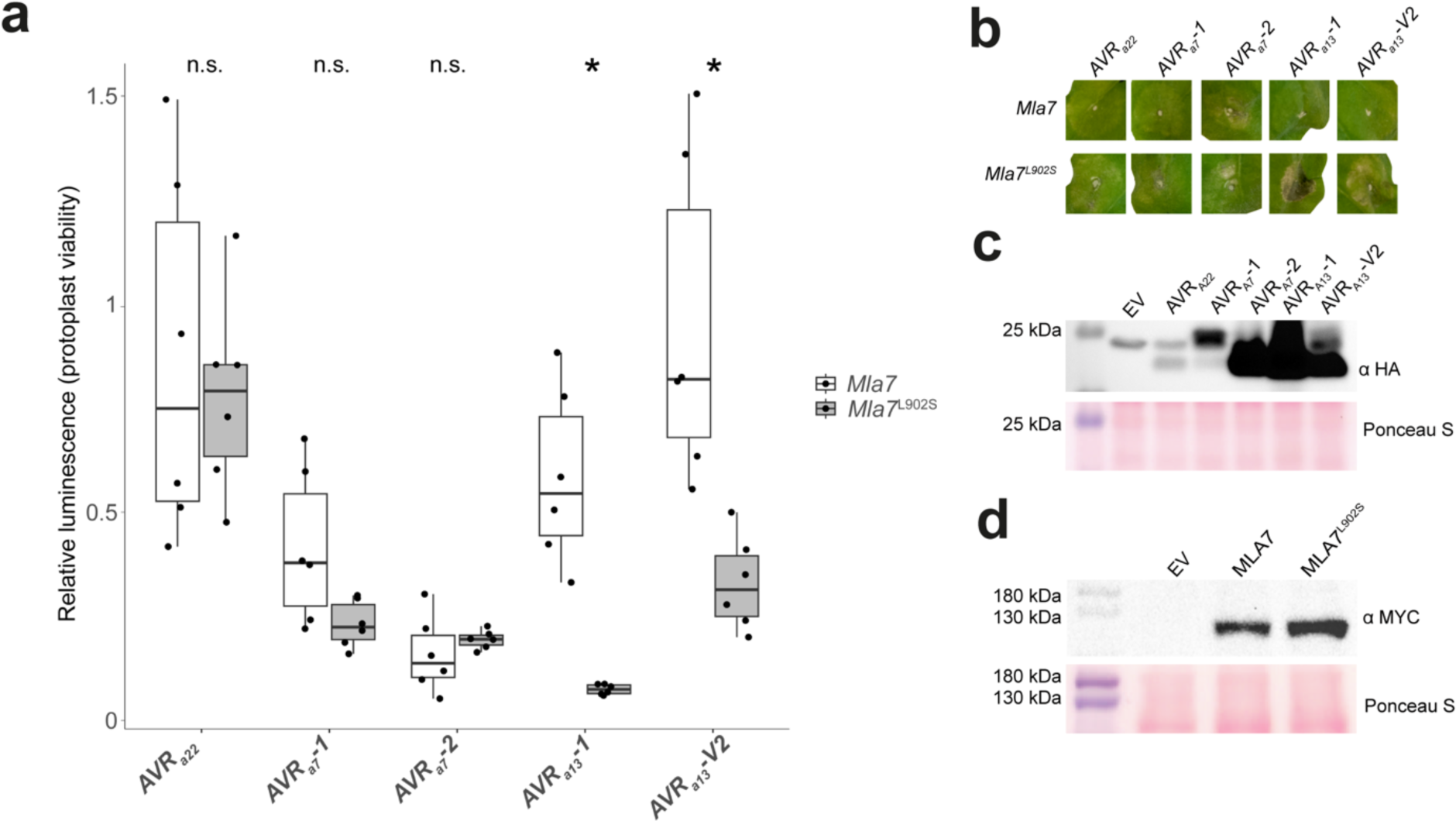
The MLA7^L902S^ substitution mutant results in expanded effector recognition. **a,** Co-expression of MLA7 and MLA7^L902S^ with AVR_A7 a_nd AVR_A13 v_ariants in barley protoplasts. Luminescence is normalised to EV + MLA7 (= 1) or EV + MLA7^L902S^ (= 1). High relative luminescence suggests low cell death response and therefore suggests low effector interaction with the receptor. The six data points represent two technical replicates performed with three independently prepared protoplast samples. Treatments labelled with an asterisk differ significantly (*p* <0.05) according to the Welch two-sample t-test. **b,** *Agrobacterium*-mediated co-expression of MLA7 and MLA7^L902S^ with AVR_A7 a_nd AVR_A13 v_ariants in *N. benthamiana* leaves. Three independent replicates were performed (Supplementary Fig. 4). **c,** Western blot analysis of the effector variants tested in (**b**). **d,** Western blot analysis of MLA7 and MLA7^L902S^.

## Discussion

Resolving the structure of the MLA13–AVR_A13_-1 heterodimer revealed a ‘noncanonical’ conformation compared to two known pentameric plant CNL resistosomes, *A. thaliana* ZAR1 and wheat Sr35^8–10^. Similar structures of monomeric ZAR1 are available (PDBs: 6J5W and 6J5V) and represent intermediate forms of the effector-activated pentameric ZAR1 resistosome^8, 30^. The ZAR1–RKS1 complex binds ADP, and subsequent PBL2^UMP^ binding in the presence of ATP results in allosteric changes, allowing the exchange of ADP to ATP in the NBD and the formation of a fully activated ZAR1 resistosome^8, 30^. ZAR1–RKS1 binding of PBL2^UMP^ in the absence of ATP results in a nucleotide-free, ligand-bound intermediate complex (PDB: 6J5V), a conformation reminiscent of the MLA13–AVR_A13_-1 heterodimer.

In contrast to the Sr35 and Sr50 resistosomes, MLA13 oligomerisation *in planta* was only detectable when introducing an autoactive-inducing substitution (MLA13^D502V^), which is thought to mimic ATP binding, resulting in effector-independent cell death (Extended Data Fig. 6). We expressed and purified a stable MLA13–AVR_A13_-1 heterodimer using the same protocol successfully used to purify pentameric Sr35 and Sr50 resistosomes. This prompts the question: why does the co-expression of MLA13 and AVR_A13_-1 not result in the purification of a higher-order complex (i.e., an MLA13 resistosome) from an *in planta* expression system (Extended Data Figs, 4,5,7)? We consider four possible explanations for this result. First, a high-order MLA13–AVR_A13_-1 heterocomplex might be prone to disassociation and thus requires yet unknown extraction conditions to maintain resistosome conformation when isolated. Second, the conformational transition between effector-dependent, intermediate and oligomeric receptor states might be differentially regulated in MLA13, Sr35, Sr50 and ZAR1. Third, in heterologous *N. benthamiana*, additional components for abundant MLA13 high-order complex formation might be present in insufficient concentrations for detectable resistosome formation. For instance, a number of MLA resistance specificities, including MLA13, require the barley co-chaperones RAR1 and SGT1 for full immunity to *Bh*^31–34^. These two proteins form a ternary HSP90–RAR1–SGT1 chaperone complex, which elevates pre-activation MLA steady-state levels in barley and might facilitate the formation of MLA13 resistosomes from the MLA13–AVR_A13_-1 heterodimer. Finally, it is also possible that the stable MLA13–AVR_A13_-1 heterodimer generates a CNL output that is distinct from CNL resistosomes. For example, it remains to be tested whether the heterodimeric complex described here contributes to nucleo-cytoplasmic partitioning of MLA receptors and their interference with the transcription machinery *via* associations with barley transcription factors^27, 35, 36^.

Structure-guided amino acid substitutions of the receptor–effector interface demonstrate the importance of MLA13–AVR_A13_-1 interactions for triggering effector-dependent and receptor-mediated plant cell death. This interface is primarily mediated by interactions supported by residues in the MLA13 WHD, LRR and two basal loops in AVR_A13_-1. Similarly, earlier structure–function analyses of AVR_A10,_ AVR_A22 a_nd AVR_A6 h_ybrid effectors suggested that multiple highly polymorphic effector surface residues in the basal loops of each of these *Bh* RALPH effectors are indispensable for recognition by their matching MLA receptors^19, 23^. This suggests the existence of a common structural principle by which functionally diversified MLA receptors recognise sequence-unrelated RALPH effectors *via* their polymorphic basal loops. This is consistent with the observation that the structural core of RALPH effectors with two β-sheets and a central α-helix of AVR_A13_-1 does not directly contribute to binding MLA13. Interestingly, Alphafold3 generated several models in which AVR_A13_-1 binds to the LRR domain of MLA13, but neither the binding site to the LRR nor the orientation of the effector relative to the LRR corresponds to the experimentally determined receptor–effector interface (Fig. 2d). Why would MLA receptors preferentially recognise AVR_A e_ffectors at the basal loops and not at other distant surface regions of the RNase-like scaffold? We hypothesise that the polymorphic sequences in the basal loops are important for the virulence activity of these *Bh* RALPH effectors, perhaps allowing them to interact with different virulence targets. However, wheat CNL Pm2a is believed to detect the *Bt* RALPH effector AvrPm2 on the opposite effector side, termed the ‘head epitope’ which comprises the juxtaposed N- and C-termini^16^. This could be explained by the finding that Pm2a recognises AvrPm2 indirectly through interaction with the wheat zinc finger protein TaZF^23^. An alternative hypothesis is that MLAs avoid recognising conserved structural elements, such as those of RNase-like scaffolds, to prevent interacting with RNase-like host proteins that may trigger a non-pathogen-induced cell death.

Here we provide evidence that residues in the C-terminal region of the MLA13 LRR are essential for receptor-mediated cell death activation upon detection of its cognate effector AVR_A13_-1. The broader relevance of the C-terminal LRR region among MLA receptors for the detection of different AVR_A e_ffectors is supported by domain swap experiments between LRR regions of MLA1 and MLA6 and MLA10 and MLA22, respectively^19, 32^. Our results show that although the LRR region is the most polymorphic among characterized MLA receptors, there are relatively few polymorphic residues in the MLA13 LRR that are critical for recognition of AVR_A13_-1^12^. This information, combined with knowledge of natural LRR sequence polymorphisms among MLA receptors with distinct AVR_A e_ffector recognition specificities, has informed the design of a GoF MLA receptor with only a single-base edit (MLA7^L902S^). Importantly, in the context of MLA13, substitution of MLA13^S902A^ resulted in a retention of AVR_A13_-1-triggered cell death activity, suggesting that MLA13^S902^ may not play a critical role in supporting the interface with AVR_a13_-1. In the context of MLA7, the MLA7^L902S^ substitution is crucial for a gain of AVR_a13_-1 detection, suggesting that the bulky MLA7^L902^ disrupts the stability of MLA7^R938^ and its essential role in effector interaction. Nevertheless, without experimental MLA7^L902S^ structures bound to AVR_A13_-1 and AVR_A7_-2, we cannot rule out the possibility that variation in the basal loop lengths of these two AVR_A e_ffectors might lead to conformationally different receptor–effector interfaces (Extended Data Figs. 9,10). In fact, the structural polymorphisms between the two RALPH subfamilies, which include AVR_A7_-2 and AVR_A13_-1, differ primarily in the lengths of the four antiparallel β-strands (β3 to β6) of the second β-sheet and not the number of structural elements, thereby resulting in different lengths of the basal loops^23^. Since the crystal structures of AVR_A6,_ AVR_A7_-2, AVR_A10,_ and AVR_A22 r_epresent unbound effector folds and a structure for unbound AVR_A13_-1 is not available, it remains to be clarified whether the basal loops of AVR_A e_ffectors undergo conformational changes upon receptor binding and, if so, whether these are similar or vary among AVR_A e_ffectors (Extended Data Figs. 9,10).

Expanding effector detection specificity by minimal perturbations such as single-base gene editing is an attractive approach for accomplishing more durable disease resistance in crops. Characterized *Mla* resistance specificities to *Bh* are alleles of one of three highly sequence-diverged CNL homologs at the complex *Mla* locus^33, 37, 38^. This precludes the generation of lines expressing two or more homozygous *Mla* resistance specificities by crossings between accessions encoding naturally polymorphic *Mla*s. The expanded detection capability of MLA7^L902S^ is a promising and notable proof-of-principle, as the receptor is able to recognise multiple RALPH effectors belonging to two phylogenetic subfamilies. The new repertoire of matching effectors detected by MLA7^L902S^ is simultaneously expressed in several globally distributed *Bgh* strains and includes the virulent effector, AVR_A13_-V2, which is presumed to be the result of resistance escape of MLA13 due to selection pressures^17, 18, 26^. Furthermore, certain allelic *Pm3* resistance specificities in wheat confer both strain-specific immunity to *Bt* and non-host resistance to other cereal mildews^14^. These wheat Pm3 CNL receptors recognise strain-specific matching *Bt* RALPH effectors and conserved RALPH effector homologues in rye mildew (*B. graminis* f sp *secale*), thereby restricting growth of rye mildew on wheat^14^. Given that barley MLA7^L902S^ also confers enhanced cell death activity to the naturally occurring virulent variant of AVR_A13_-1, AVR_A13_-V2, and that the 34 members of this RALPH subfamily include several *Bt* effectors, including AvrPm2 and *Bt* E-5843, it seems possible that this or other engineered MLA receptors could enhance barley non-host resistance to other cereal mildews^16, 17, 23^. Future work will complement our findings by generating gene edited barley lines expressing synthetic MLAs for resistance testing.

## Methods

### Plant growth

Seeds of wild-type *N. benthamiana* were sown in peat-based potting soil with granulated cork on the surface to prevent pest infestation. Daily irrigation solution contained an electrical conductivity of 2.2 and a mixture of macro and micro nutrients. A photoperiod of 16 hours was used with broad-spectrum LED lights emitting 220 μmol/m^2^/s supplemented by ambient sunlight.

Barley protoplasts isolated from Golden Promise seedlings that were grown on peat-based potting soil at 19 °C and 70% humidity for 7–9 days.

### Transient transformation of *N. benthamiana* for recombinant protein expression and purification

The coding sequences of *Mla13* containing a stop codon was transferred from pDONR221 using Gateway LR clonase into pGWB424 containing an N-terminal fusion GST tag in the vector backbone. *AVR_a13_-1* without a stop codon was transferred from pDONR221 using Gateway LR clonase into pGWB402SC containing a C-terminal Twin-Strep-tag® followed by a single HA tag in the vector backbone. Both constructs were individually electroporated into *Agrobacterium tumefaciens* strain GV3101::pMP90RK and selected on plates of Luria/Miller (LB) broth with agar containing spectinomycin (100 μg/mL), gentamycin (25 μg/mL), rifampicin (50 μg/mL) and kanamycin (25 μg/mL) and grown for two days at 28 °C. Three colonies were picked and cultured overnight in a 10-mL liquid LB starter culture with the above antibiotics at 28 °C. Two millilitres of the starter culture were added to and cultured in 350 mL of liquid LB broth containing the above antibiotics for 14 hours at 28 °C. The cultures were pelleted at 4,000 RCF for 15 minutes and resuspended in infiltration buffer (10 mM MES (pH 5.6), 10 mM MgCl_2,_ 500 μM acetosyringone) to an OD_600 o_f 2 for each construct. The bacterial suspensions were combined at a 1:1 ratio and infiltrated into leaves of four-week-old *N. benthamiana* plants. The infiltrated plants were stored in the dark for 24 hours before they were returned to normal growth conditions where they grew for an additional 24 hours. The leaves were frozen in liquid nitrogen and stored at −80 °C until they were processed.

### Protein purification for cryo-EM

One hundred grams of transiently transformed *N. benthamiana* leaf tissue were ground in a prefrozen mortar and pestle and gradually added to 200 mL of lysis buffer (buffer A; 50 mM Tris-HCl (pH 7.4), 150 mM NaCl, 5% glycerol, 10 mM DTT, 0.5% polysorbate 20, two vials of protease inhibitor cocktail (SERVA Electrophoresis GmbH catalogue # 39103.03), 5% BioLock (IBA Lifesciences GmbH catalogue # 2-02-5-250); pH adjusted to 7.4) until the lysate was defrosted and at 4 °C. The lysate was split into two 250 mL centrifuge bottles, centrifuged twice at 30,000 RCF for 15 minutes and filtered through double-layered miracloth after each centrifuge run.

Five hundred microlitres of Strep-Tactin XT Sepharose resin (Cytiva catalogue # 29401324) were equilibrated in wash buffer (buffer B; 50 mM Tris-HCl (pH 7.4), 150 mM NaCl, 2 mM DTT, 0.1% polysorbate 20; pH adjusted to 7.4). The resin was added to the lysate and incubated by end-over-end rotation at 4 °C for 30 minutes. The resin was washed three times with buffer B and finally isolated in a 1.5-mL tube. Five hundred microlitres of Strep-Tactin XT Sepharose resin elution buffer (buffer C; buffer B supplemented with 50 mM biotin; pH adjusted to 7.4) was added to the resin and rotated end-over-end for 30 minutes. The above elution step was repeated five times.

The five eluates were centrifuged at 16,000 RCF for one minute and 450 μL of supernatant were removed from each eluate and pooled. Two hundred microlitres of Glutathione Sepharose 4B resin (Cytiva catalogue # 17075601) was equilibrated in buffer B and added to the Strep-Tactin XT eluate was combined with the Glutathione Sepharose 4B resin and incubated by mixing end-over-end for two hours at 4 °C. The Glutathione Sepharose 4B resin was washed twice before with buffer B. Elution from the Glutathione Sepharose 4B resin was performed by adding 200 μL of buffer D (buffer B supplemented with 50 mM reduced glutathione; pH adjusted to 7.4) and rotated end-over-end for 30 minutes. Elution was repeated for a total of four times. The four eluates were centrifuged at 16,000 RCF for one minute and 150 μL of supernatant were removed from each eluate. Twenty microlitres from the first eluate were used for cryo-EM grid preparation and the remaining eluate(s) were pooled and analysed by SEC.

For SEC, a Superose 6 increase 10/300 GL column (Cytiva catalogue #) was equilibrated with buffer B. Five hundred microlitres of the pooled GST eluate were loaded into the column and run at 0.3 mL/minute. Forty-five microlitres of the 500 μL fractions were loaded on SDS PAGE gels.

The Sr35 and Sr50 resistosomes were purified with the above method. The *in planta* cell death activity was abrogated for purification purposes through introduction of the L11E/L15E substitutions in the receptors. A single-step purification was performed by coimmunoprecipitating the effectors *via* the C-terminal and N-terminal twin-Strep epitope tags on AvrSr35 and AvrSr50, respectively. Sr35 and Sr50 were expressed without an epitope tag. The 5 mL of twin-Strep eluate was concentrated and analysed by SEC as described above.

### Negative staining and TEM

Carbon film grids (Electron Microscopy Sciences catalogue # CF400-CU-50) were glow discharged for negative staining of protein samples. The MLA13-AVR_A13_-1 heterodimer, Sr35 resistosome and Sr50 resistosome samples were series-diluted in buffer B. Six microlitres of sample were applied to the grid and incubated for one minute before blotting off excess sample with filter paper. Six microlitres of one percent uranyl acetate were then applied to the grids and incubated for one minute before blotting off with filter paper.

Grids were analysed using a Hitachi HT7800 TEM operating at 100 kV and fitted with an EMSIS XAROSA camera.

### Cryo-EM sample preparation and data collection

Three microlitres of the purified MLA13–AVR_A13_-1 sample were applied to an untreated graphene oxide-coated TEM grid (Science Services catalogue # ERGOQ200R24Cu50), incubated on the grid for 10 seconds, blotted for 5 seconds and flash-frozen in liquid ethane using a Vitrobot Mark IV device (Thermo Fisher Scientific) set to 90% humidity at 4 °C. Grids were stored under liquid nitrogen conditions until usage.

Cryo-EM data was acquired using a Titan Krios G3i (Thermo Fisher Scientific) electron microscope operated at 300 kV. Images were collected automatically using EPU (version 2.12) (Thermo Fisher Scientific) on a Falcon III direct electron detector with a calibrated pixel size of 0.862 Å*px^−1^. Target defocus values were set to −2.0 to −0.3 μm. Data was acquired using a total dose of 42 e^−^*Å^−2^ distributed among 42 frames, although the last three frames were excluded during data analysis.

### Image processing and model building

Image processing was performed using CryoSPARC (version 4.1.1+patch 240110). Movie stacks were first corrected for drift and beam-induction motion, and then used to determine defocus and other CTF-related values. Only high-quality micrographs with low drift metrics, low astigmatism, and good agreement between experimental and calculated CTFs were further processed. Putative particles were automatically picked based on an expected protein diameter between 8 and 12 nm, then extracted and subjected to reference-free 2D classification. 2D classes showing protein-like shapes were used for a template-based picking approach. Candidate particles were extracted again, subjected to reference-free 2D classification to exclude artefacts, and subsequent 3D classification to identify high-quality particles showing defined density for the effector, NBD, and LRR. This subset of particles was further refined using the non-uniform refinement strategy, yielding a map at a global resolution of 3.8 Å. DeepEMhancer was used to optimize the map for subsequent structure building. For further details see Extended Data Fig.2.

AlphaFold was used to predict a model for the CC-NBD-LRR domains of MLA13 from *H. vulgare* using the sequence Q8GSK4 from UniProt and two previously deposited structures in the PDBe, 5T1Y and 3QFL. The AlphaFold-predicted model was fitted into the map; however, the fold of the CC-domain did not match the observed density adjacent to the LRR. Afterward, Robetta was used to predict only this region, which gave outputs that more closely resembled the activated form of ZAR1 resistosome’s CC-domain. Robetta uses deep learning-based methods, RoseTTAFold and TrRosetta algorithms, and thus it may be influenced by existing models of the sequence to be predicted. For this reason, the *ab initio* option was chosen when running a second round of predictions in Robetta, and a template of the inactive ZAR1 CC-domain from *A. thaliana* (6J5W, Wang et al 2019) was included in the subsequent prediction run. The new model of the CC-domain fitted the EM map significantly better than the previous predicted models; thus, it was merged with the rest of the MLA13 model for refinement. Finally, the model containing AVR_A13_-1-bound MLA13 was refined against the EM map in iterations of *phenix.real_space_refine* and manual building in Coot. For further details and statistics see Supplementary Table 1. Molecular visualization and analysis were done using UCSF ChimeraX (version 1.7).

### Cell death assays in barley protoplasts

Experiments were performed according to Saur *et al.* 2019 with the exception that plasmid DNA of all constructs was diluted to 500 ng/μL and transfection volumes were 15 μL, 10 μL, and 10 μL for *pUBQ:luciferase*, *Mla*, and *AVR_a_*, respectively^39^.

### Cell death assays in leaves of *N. benthamiana*

DNA of effector and receptor sequences were cloned as mentioned above into pGWB402SC and pGWB517, respectively. Transformation and preparation of *A. tumefaciens* suspensions was performed as mentioned above. Phenotype images were taken 72 hours post infiltration while samples for western blot analysis were harvested 24 hours post infiltration.

Western blotting of samples consisted of flash-freezing 100 mg of each sample and pulverising the tissue using a bead beater. The frozen leaf powder was resuspended in the aforementioned buffer A. The samples were centrifuged twice at 16,000 RCF before adding 4× Lämmli buffer (Bio-Rad catalogue # 161-0737) supplemented with 5% mM β-mercaptoethanol and heating the sample to 95 °C for five minutes before cooling on ice. Ten microlitres of each sample were run on 12% SDS PAGE gels before transferring to a PVDF membrane. The membranes were then blocked in TBS-T containing 5% milk for one hour at room temperature (RT). Membranes were washed three times for five minutes in TBS-T then incubated with anti-HA (Cell Signalling Technology catalogue # 3724; 1:1,000) and anti-MYC (Thermo Fisher Scientific Inc. catalogue # R950-25; 1:5,000) in TBS-T with 5% BSA for one hour at RT. Membranes were washed in TBS-T for 3 × 10 minutes incubating with secondary anti-rabbit (Cell Signalling Technology catalogue # 7074S; 1:2,000) and anti-mouse (Abcam Ltd. Catalogue # ab6728; 1:5,000) in TBS-T with 5% milk for one hour at RT. Membranes were washed in TBS-T for 3 × 15 minutes before developing using SuperSignal West Femto substrate (Thermo Fisher Scientific Inc. catalogue # 34096).

### BN-PAGE assays

BN-PAGE assays were performed as described in Ma *et al.* (2024) with modifications^40^. Briefly, *N. benthamiana* leaf tissues expressing the indicated constructs were harvested at 48 h after infiltration. Two grams of each sample were ground into powder using liquid nitrogen and homogenized in 4-mL protein extraction buffer (10% glycerol, 50 mM Tris-HCl (pH 7.5), 150 mM NaCl, 5 mM DTT, 0.2% NP-40, 5 mM MgCl_2,_ 20 µM MG132, 1×Roche protease inhibitor cocktail). The extract was centrifuged twice at 4 °C, 12,000 RCF for 15 min. Then, 40 µL of extraction buffer-washed Strep-Tactin® Sepharose chromatography resin (Cytiva) were added to the extract and incubated with end-over-end rotation for one hour. The resins were collected by centrifugation at 1,000 RCF for 3 min and washed three times with wash buffer (10% glycerol, 50 mM Tris-HCl (pH 7.5), 50 mM NaCl, 2 mM DTT, 0.2% NP-40, 1×Roche protease inhibitor cocktail). Subsequently, 100 µL of elution buffer (wash buffer + 50 mM biotin) were added to the resin and followed by end-over-end rotation for 30 min. The purified protein samples were collected by centrifugation. Five microlitres of each sample (25 µL for MLA13 auto-active mutants) were mixed with Native PAGE G-250 additive to a final concentration of 0.1%, and placed on ice for 30 min. Protein samples and unstained Native Mark (Invitrogen catalogue #LC0725) were loaded and run on a Native PAGE 3%-12% Bis-Tris gel (Invitrogen catalogue #BN1001BOX) according to the manufacturer’s instructions. The proteins were then immunoblotted as described above.

## Data availability

The EM map has been deposited in the EMDB under the accession code EMD-50863. Atomic coordinates have been deposited in the Protein Data Bank under the accession code 9FYC. Other data used to generate tables and figures has been provided as source data with this publication.

## Acknowledgements

We thank the greenhouse team at MPIPZ for their expertise in providing high-quality *N. benthamiana* plants. We thank Neysan Donnelly and Jane Parker for critical comments on an early version of this manuscript. We thank Arthur Macha, Petra Koechner, Sabine Haigis, Elke Logemann, Milena Malisic, Florian Kuemmel, Li Liu, Wen Song and Nitika Mukhi for their intellectual and experimental contributions. This work was funded by the Max-Planck-Gesellschaft (P.S.-L.), the Deutsche Forschungsgemeinschaft (DFG, German Research Foundation) in the Collaborative Research Centre Grant (SFB-1403 – 414786233 B08 to P.S.-L., I.M.L.S. and J.C.), Germany’s Excellence Strategy CEPLAS (EXC-2048/1, project 390686111 to P.S.-L. and I.M.L.S.), the Ministry of Culture and Science of the State of North Rhine-Westphalia (iHEAD to P.S.-L. and E.B.) and DFG Emmy Noether Programme (SA 4093/1-1 to I.M.L.S.). We acknowledge access to the cryo-EM infrastructure of StruBiTEM (Cologne, funded by DFG Grant INST 216/949-1 FUGG), and to the computing infrastructure of CHEOPS (Cologne, funded by DFG Grant INST 216/512/1 FUGG).

## Author contributions

P.S.-L., J.C., E.B. and A.W.L. conceived the study; A.W.L., Y.C., C.A., M.G. and I.M.L.S. performed experiments; U.N. and M.G. performed electron microscopy screening; A.W.L., E.B., J.C. and P.S.-L. analysed data; A.F.-I. and E.B. performed structural model building; P.S.-L., E.B. and A.W.L. wrote the manuscript.

## References

1. S. T. Chisholm, G. Coaker, B. Day and B. J. Staskawicz, Cell 124 (4), 803–814 (2006).

2. T. Maekawa, B. Kracher, S. Vernaldi, E. Ver Loren van Themaat and P. Schulze-Lefert, Proceedings of the National Academy of Sciences 109 (49), 20119–20123 (2012).

3. J. Chai, W. Song and J. E. Parker, Molecular Plant-Microbe Interactions® 36 (8), 468–475 (2023).

4. I. M. L. Saur, R. Panstruga and P. Schulze-Lefert, Nature Reviews Immunology 21 (5), 305–318 (2020).

5. S. M. Collier, L.-P. Hamel and P. Moffett, MPMI 24 (8), 918–931 (2011).

6. A. Bentham, H. Burdett, P. A. Anderson, S. J. Williams and B. Kobe, Annals of Botany (2016).

7. J. D. G. Jones, B. J. Staskawicz and J. L. Dangl, Cell 187 (9), 2095–2116 (2024).

8. J. Wang, M. Hu, J. Wang, J. Qi, Z. Han, G. Wang, Y. Qi, H.-W. Wang, J.-M. Zhou and J. Chai, Science 364 (6435) (2019).

9. A. Förderer, E. Li, A. W. Lawson, Y. N. Deng, Y. Sun, E. Logemann, X. Zhang, J. Wen, Z. Han, J. Chang, Y. Chen, P. Schulze-Lefert and J. Chai, Nature 610 (7932), 532–539 (2022).

10. Y. B. Zhao, M. X. Liu, T. T. Chen, X. Ma, Z. K. Li, Z. Zheng, S. R. Zheng, L. Chen, Y. Z. Li, L. R. Tang, Q. Chen, P. Wang and S. Ouyang, Sci Adv 8 (36), eabq5108 (2022).

11. G. Bi, M. Su, N. Li, Y. Liang, S. Dang, J. Xu, M. Hu, J. Wang, M. Zou, Y. Deng, Q. Li, S. Huang, J. Li, J. Chai, K. He, Y.-h. Chen and J.-M. Zhou, Cell 184 (13), 3528–3541.e3512 (2021).

12. S. Seeholzer, T. Tsuchimatsu, T. Jordan, S. Bieri, S. Pajonk, W. Yang, A. Jahoor, K. K. Shimizu, B. Keller and P. Schulze-Lefert, Mol Plant Microbe Interact 23 (4), 497–509 (2010).

13. S. Bourras, K. E. McNally, R. Ben-David, F. Parlange, S. Roffler, C. R. Praz, S. Oberhaensli, F. Menardo, D. Stirnweis, Z. Frenkel, L. K. Schaefer, S. Flückiger, G. Treier, G. Herren, A. B. Korol, T. Wicker and B. Keller, Plant Cell 27 (10), 2991–3012 (2015).

14. S. Bourras, L. Kunz, M. Xue, C. R. Praz, M. C. Müller, C. Kälin, M. Schläfli, P. Ackermann, S. Flückiger, F. Parlange, F. Menardo, L. K. Schaefer, R. Ben-David, S. Roffler, S. Oberhaensli, V. Widrig, S. Lindner, J. Isaksson, T. Wicker, D. Yu and B. Keller, Nature Communications 10 (1), 2292 (2019).

15. C. R. Praz, S. Bourras, F. Zeng, J. Sánchez-Martín, F. Menardo, M. Xue, L. Yang, S. Roffler, R. Böni, G. Herren, K. E. McNally, R. Ben-David, F. Parlange, S. Oberhaensli, S. Flückiger, L. K. Schäfer, T. Wicker, D. Yu and B. Keller, New Phytol 213 (3), 1301–1314 (2017).

16. B. Manser, T. Koller, C. R. Praz, A. C. Roulin, H. Zbinden, S. Arora, B. Steuernagel, B. B. H. Wulff, B. Keller and J. Sánchez-Martín, Plant J 106 (4), 993–1007 (2021).

17. X. Lu, B. Kracher, I. M. L. Saur, S. Bauer, S. R. Ellwood, R. Wise, T. Yaeno, T. Maekawa and P. Schulze-Lefert, Proceedings of the National Academy of Sciences 113 (42) (2016).

18. I. M. L. Saur, S. Bauer, B. Kracher, X. Lu, L. Franzeskakis, M. C. Müller, B. Sabelleck, F. Kümmel, R. Panstruga, T. Maekawa and P. Schulze-Lefert, eLife 8 (2019).

19. S. Bauer, D. Yu, A. W. Lawson, I. M. L. Saur, L. Frantzeskakis, B. Kracher, E. Logemann, J. Chai, T. Maekawa and P. Schulze-Lefert, PLOS Pathogens 17 (2) (2021).

20. C. Pedersen, E. Ver Loren van Themaat, L. J. McGuffin, J. C. Abbott, T. A. Burgis, G. Barton, L. V. Bindschedler, X. Lu, T. Maekawa, R. Wessling, R. Cramer, H. Thordal-Christensen, R. Panstruga and P. D. Spanu, BMC Genomics 13, 694 (2012).

21. H. G. Pennington, R. Jones, S. Kwon, G. Bonciani, H. Thieron, T. Chandler, P. Luong, S. N. Morgan, M. Przydacz, T. Bozkurt, S. Bowden, M. Craze, E. J. Wallington, J. Garnett, M. Kwaaitaal, R. Panstruga, E. Cota and P. D. Spanu, PLOS Pathogens 15 (3), e1007620 (2019).

22. K. Seong and K. V. Krasileva, Nat Microbiol 8 (1), 174–187 (2023).

23. Y. Cao, F. Kümmel, E. Logemann, J. M. Gebauer, A. W. Lawson, D. Yu, M. Uthoff, B. Keller, J. Jirschitzka, U. Baumann, K. Tsuda, J. Chai and P. Schulze-Lefert, Proceedings of the National Academy of Sciences 120 (2023).

24. S. Bai, J. Liu, C. Chang, L. Zhang, T. Maekawa, Q. Wang, W. Xiao, Y. Liu, J. Chai, F. L. Takken, P. Schulze-Lefert and Q. H. Shen, PLoS Pathog 8 (6), e1002752 (2012).

25. T. Maekawa, W. Cheng, L. N. Spiridon, A. Töller, E. Lukasik, Y. Saijo, P. Liu, Q. H. Shen, M. A. Micluta, I. E. Somssich, F. L. W. Takken, A. J. Petrescu, J. Chai and P. Schulze-Lefert, Cell Host Microbe 9 (3), 187–199 (2011).

26. E. E. Crean, M. Bilstein-Schloemer, T. Maekawa, P. Schulze-Lefert, I. M. L. Saur and W.- M. Wang, Journal of Experimental Botany 74 (18), 5854–5869 (2023).

27. B. Manser, H. Zbinden, G. Herren, J. Steger, J. Isaksson, S. Bräunlich, T. Wicker and B. Keller, Plant Commun 5 (5), 100769 (2024).

28. Z. Wang, X. Liu, J. Yu, S. Yin, W. Cai, N. H. Kim, F. El Kasmi, J. L. Dangl and L. Wan, Proceedings of the National Academy of Sciences 120 (32) (2023).

29. J. Chen, N. M. Upadhyaya, D. Ortiz, J. Sperschneider, F. Li, C. Bouton, S. Breen, C. Dong, B. Xu, X. Zhang, R. Mago, K. Newell, X. Xia, M. Bernoux, J. M. Taylor, B. Steffenson, Y. Jin, P. Zhang, K. Kanyuka, M. Figueroa, J. G. Ellis, R. F. Park and P. N. Dodds, Science 358 (6370), 1607–1610 (2017).

30. J. Wang, J. Wang, M. Hu, S. Wu, J. Qi, G. Wang, Z. Han, Y. Qi, N. Gao, H.-W. Wang, J.-M. Zhou and J. Chai, Science 364 (6435) (2019).

31. S. Bieri, S. Mauch, Q.-H. Shen, J. Peart, A. Devoto, C. Casais, F. Ceron, S. Schulze, H.-H. Steinbiß, K. Shirasu and P. Schulze-Lefert, The Plant Cell 16 (12), 3480–3495 (2004).

32. Q.-H. Shen, F. Zhou, S. Bieri, T. Haizel, K. Shirasu and P. Schulze-Lefert, The Plant Cell 15 (3), 732–744 (2003).

33. D. A. Halterman and R. P. Wise, The Plant Journal 38 (2), 215–226 (2004).

34. A. V. E. Chapman, M. Hunt, P. Surana, V. Velásquez-Zapata, W. Xu, G. Fuerst and R. P. Wise, Genetics 217 (2) (2020).

35. Q. H. Shen, Y. Saijo, S. Mauch, C. Biskup, S. Bieri, B. Keller, H. Seki, B. Ulker, I. E. Somssich and P. Schulze-Lefert, Science 315 (5815), 1098–1103 (2007).

36. C. Chang, D. Yu, J. Jiao, S. Jing, P. Schulze-Lefert and Q. H. Shen, Plant Cell 25 (3), 1158–1173 (2013).

37. F. Wei, R. A. Wing and R. P. Wise, Plant Cell 14 (8), 1903–1917 (2002).

38. T. Maekawa, B. Kracher, I. M. L. Saur, M. Yoshikawa-Maekawa, R. Kellner, A. Pankin, M. von Korff and P. Schulze-Lefert, Mol Plant Microbe Interact 32 (1), 107–119 (2019).

39. I. M. L. Saur, S. Bauer, X. Lu and P. Schulze-Lefert, Plant Methods 15 (1) (2019).

40. S. Ma, C. An, A. W. Lawson, Y. Cao, Y. Sun, E. Y. J. Tan, J. Pan, J. Jirschitzka, F. Kümmel, N. Mukhi, Z. Han, S. Feng, B. Wu, P. Schulze-Lefert and J. Chai, Nature (2024).

