## Extended and Supplementary Figures for "The barley MLA13-AVR_A13_ heterodimer reveals principles for immunoreceptor recognition of RNase-like powdery mildew effectors"

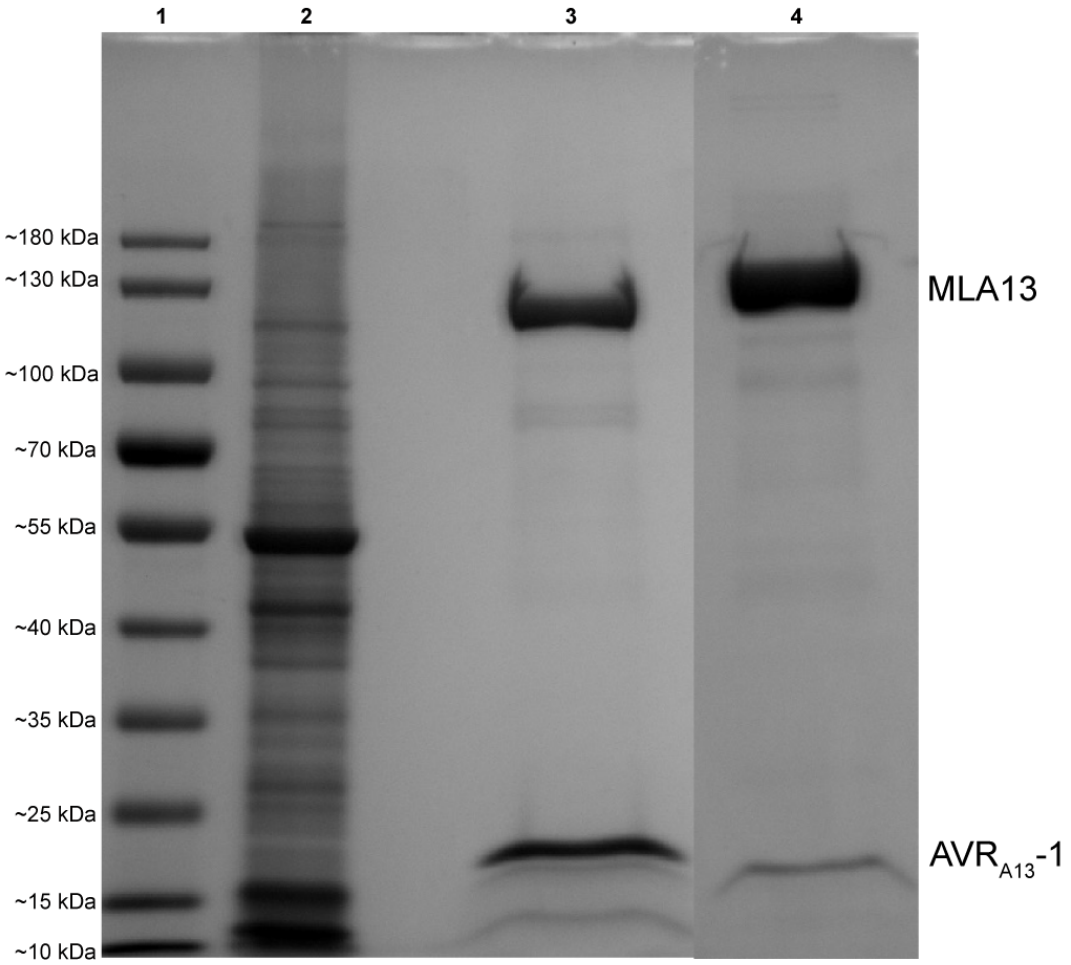


**Extended Data Fig. 1 | CBB-stained SDS PAGE gel containing the samples from a two-step affinity purification of the MLA13-AVR$_{A13}$-1 heterodimer.** Lane #1: ladder; lane #2: lysate (5 μL loaded); lane #3: first-step Twin-Strep elution (45 μL/1 mL loaded); lane #4: second-step GST elution (45 μL/750 μL loaded).

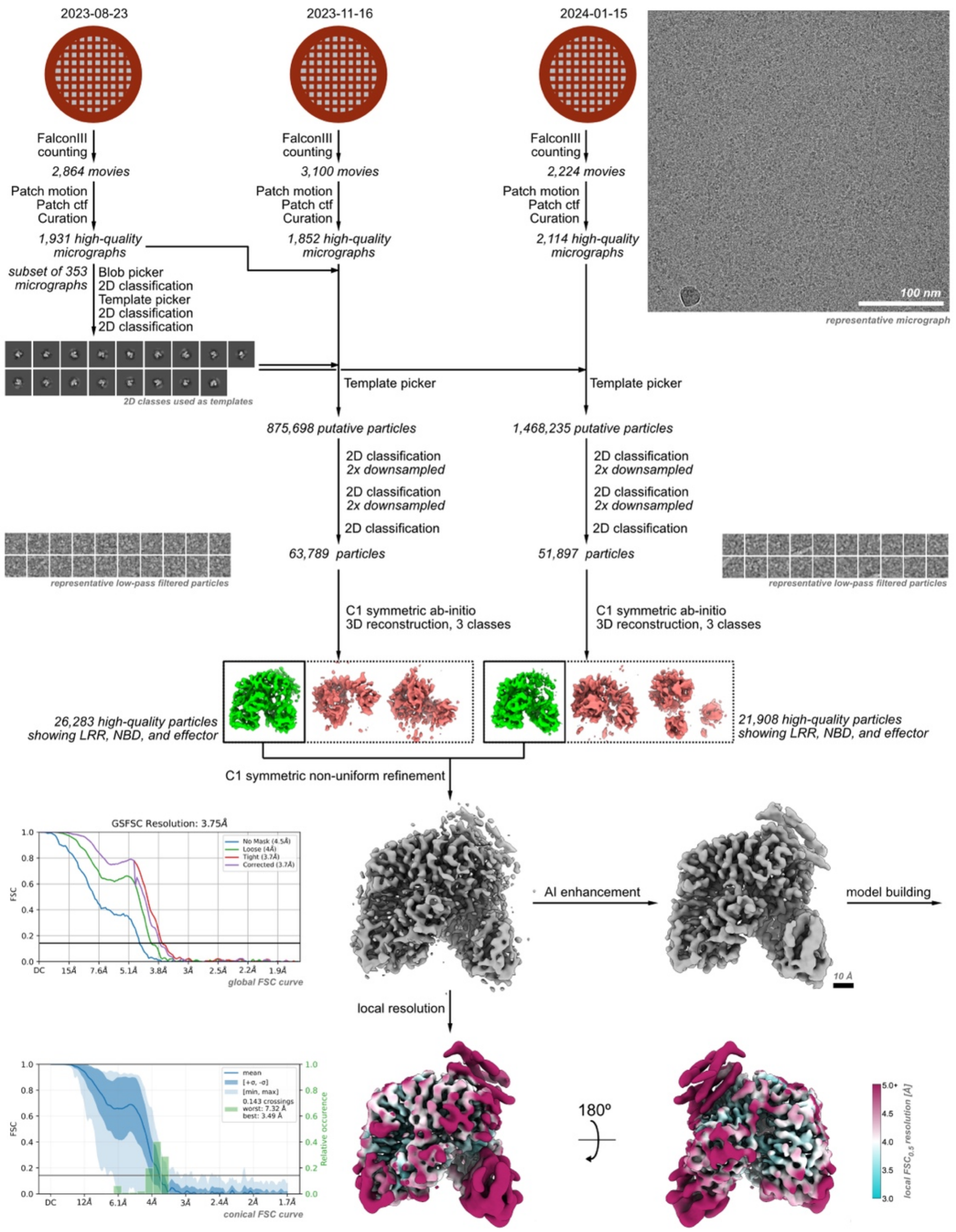


**Extended Data Fig. 2 | Workflow of cryo-EM data acquisition and analysis of the MLA13-AVR$_{A13}$-1 heterodimer.** A total of three datasets were collected on a 300 kV cryo-electron microscope. For each dataset, movies were selected for low per-frame drift rates, good CTF scores, and low astigmatism. Particles were first picked using a blob picker, and then subjected to unsupervised 2D classification. Representative classes showing protein-like structures were used for a template picker. Putative detected particles were curated using unsupervised 2D classification, selecting for particles with protein-like density and resolutions better than 10 Å. The selected particles were further curated using *ab initio* reconstruction, sorting them into three distinct populations. From these, all particles contributing to a structure showing clear density for the LRR, NBD and effector (shown in green and highlighted by a thicker box outline) were combined and refined in 3D using a non-uniform refinement algorithm, resulting in a map with a uniform resolution of 2.8 Å. Before model building, the map was further sharpened using DeepEMhancer. For further details, see the Materials and Methods and Supplementary Table 1.

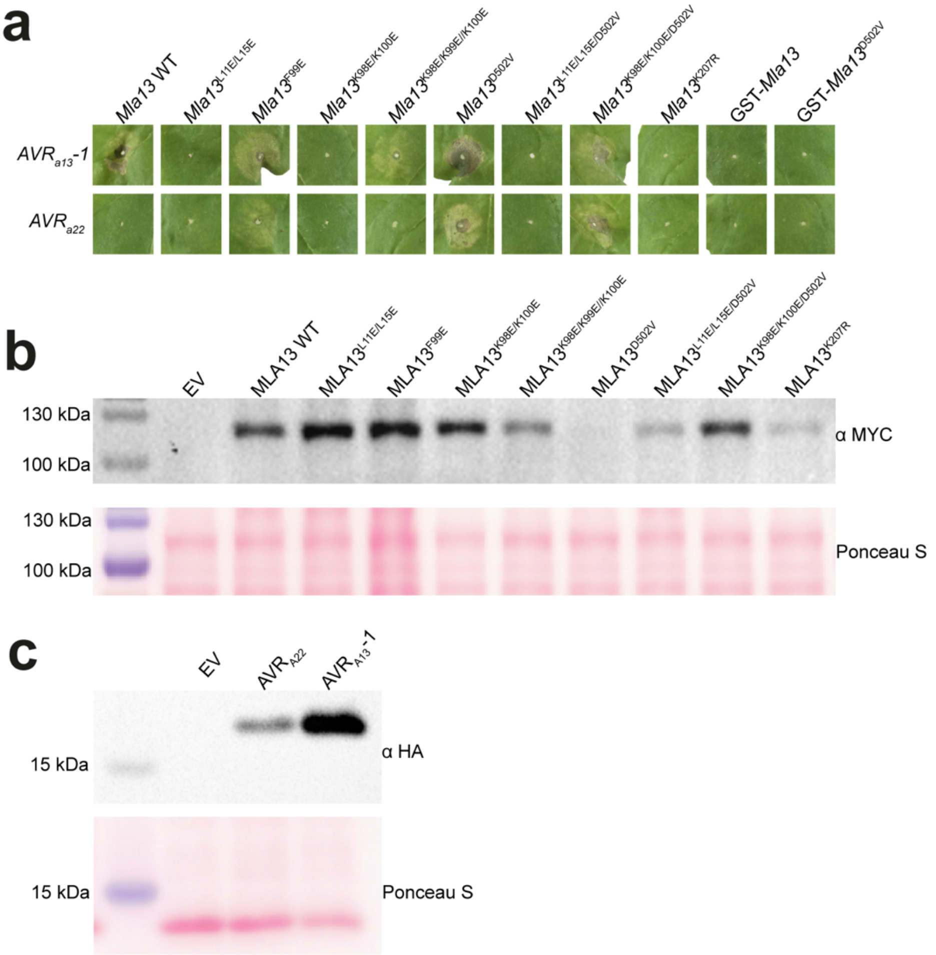


**Extended Data Fig. 3 | *Agrobacterium*-mediated co-expression of MLA13 variants with $AVR_{A13}$-1 and $AVR_{A22}$ in leaves of *N. benthamiana*. a,** Cell death phenotypes of MLA13 phenotypes of variants that result in effector-triggered HR, autoactive HR or loss of HR. Six independent replicates were performed (Supplementary Fig.1). **b,** Western blot analysis of the above MLA13 variants. **c,** Western blot analysis of $AVR_{A13}$-1 and $AVR_{A22}$.

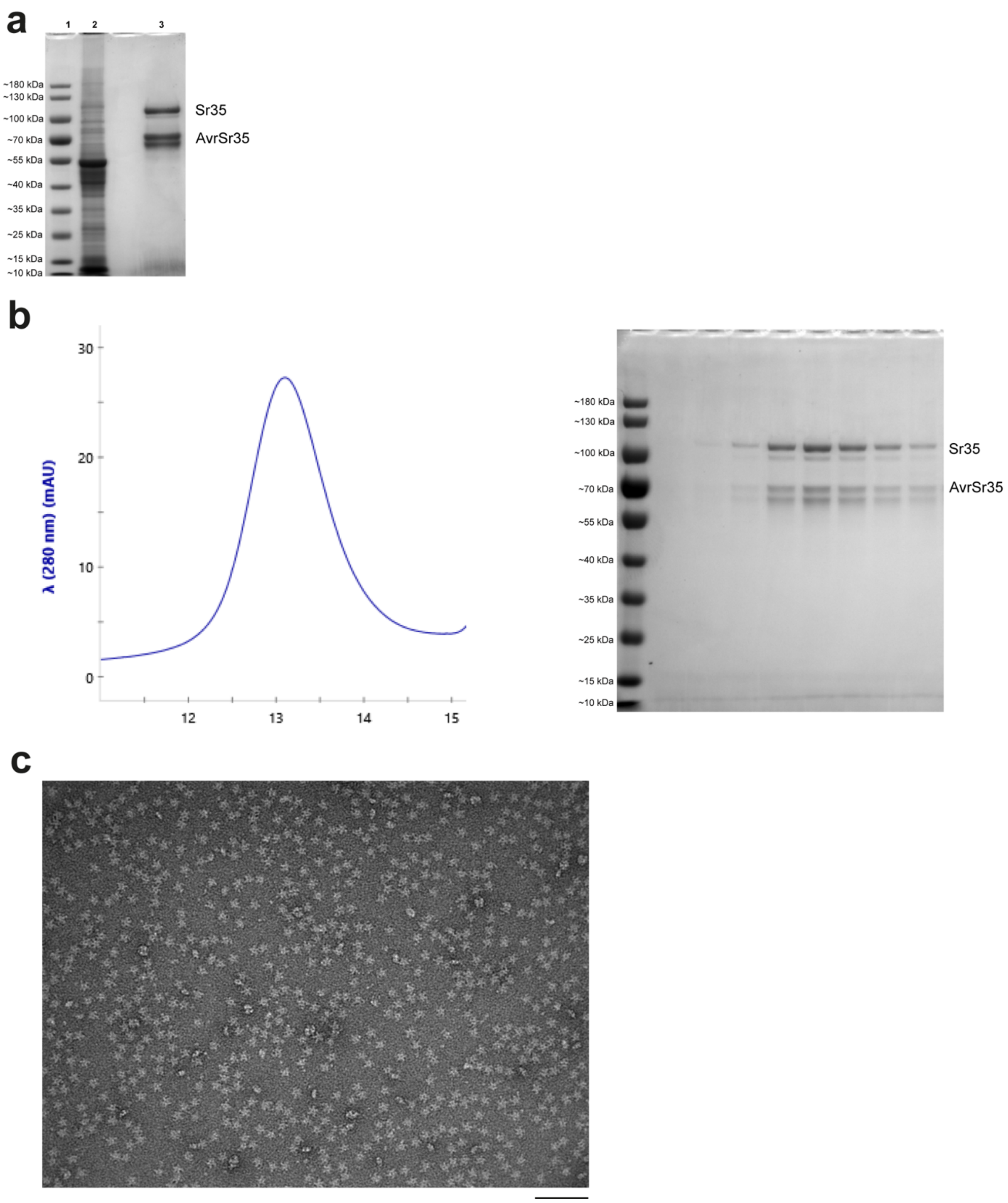


**Extended Data Fig. 4 | Transient expression and purification of the Sr35 resistosome from leaves of *N. benthamiana*. a,** CBB-stained SDS PAGE gel of a single step affinity purification of the Sr35 resistosome *via* the Twin-Strep tag at the C-terminal of AvrSr35 (Lane #1: ladder; lane #2: lysate (5 μL loaded); lane #3: first-step elution (45 μL/1 mL loaded). **b,** SEC profile (left) and SDS PAGE of resulting elution fractions. **c,** Negative staining of a five-fold dilution from the fraction corresponding to the 13 mL elution volume in (**b**). Black scale bar at the bottom right represents 100 nm.

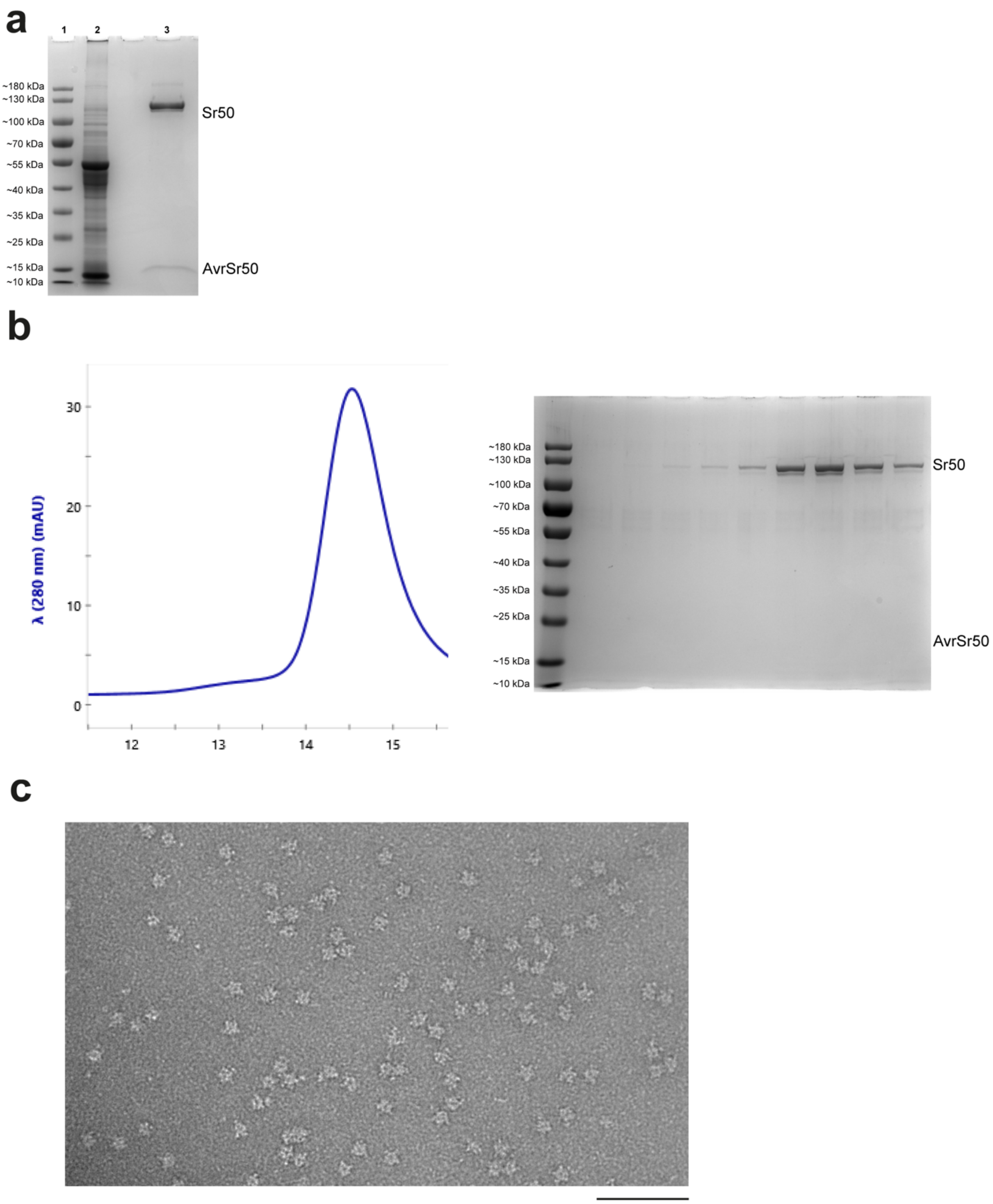


**Extended Data Fig. 5 | Transient expression and purification of the Sr50 resistosome from leaves of *N. benthamiana*. a,** CBB-stained SDS PAGE gel of a single step affinity purification of the Sr50 resistosome *via* the Twin-Strep tag at the N-terminal of AvrSr50 (Lane #1: ladder; lane #2: lysate (5 μL loaded); lane #3: first-step elution (45 μL/1 mL loaded). **b,** SEC profile (left) and SDS PAGE of resulting elution fractions. **c,** Negative staining of a 5× dilution from the fraction corresponding to the 14.5 mL elution volume in (**b**). Black scale bar at the bottom right represents 100 nm.

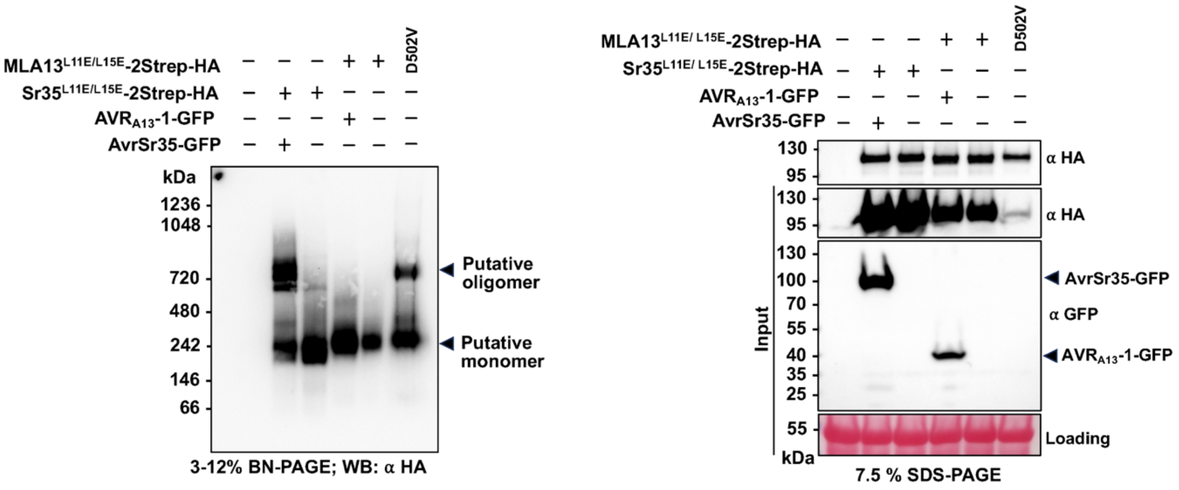


**Extended Data Fig. 6 | Blue native-PAGE analysis of MLA13 oligomeric status.** C terminal twin Strep-HA-tagged MLA13$^{L11E/L15E}$ or Sr35$^{L11E/L15E}$ were co-expressed with or without C terminally-GFP-tagged matching effectors in *N. benthamiana* leaves. The substitutions in MLA13$^{L11E/L15E}$ and Sr35$^{L11E/L15E}$ were introduced to prevent cell death. Purified protein samples *via* the twin Strep tag were analysed by blue native-PAGE (left panel) with subsequent western blotting. Deduced low-molecular weight receptor complexes, receptor monomers and oligomers are indicated by arrows. SDS-PAGE analysis of the input samples (right panel) was conducted to validate the expression of input proteins. Due to low abundance of MLA13$^{L11E/L15E/D520V}$, we loaded five times the volume of this sample on the blue native-PAGE gel. Ponceau S staining of RuBisCO was used as a loading control (right panel). Two independent experiments were performed with similar results.

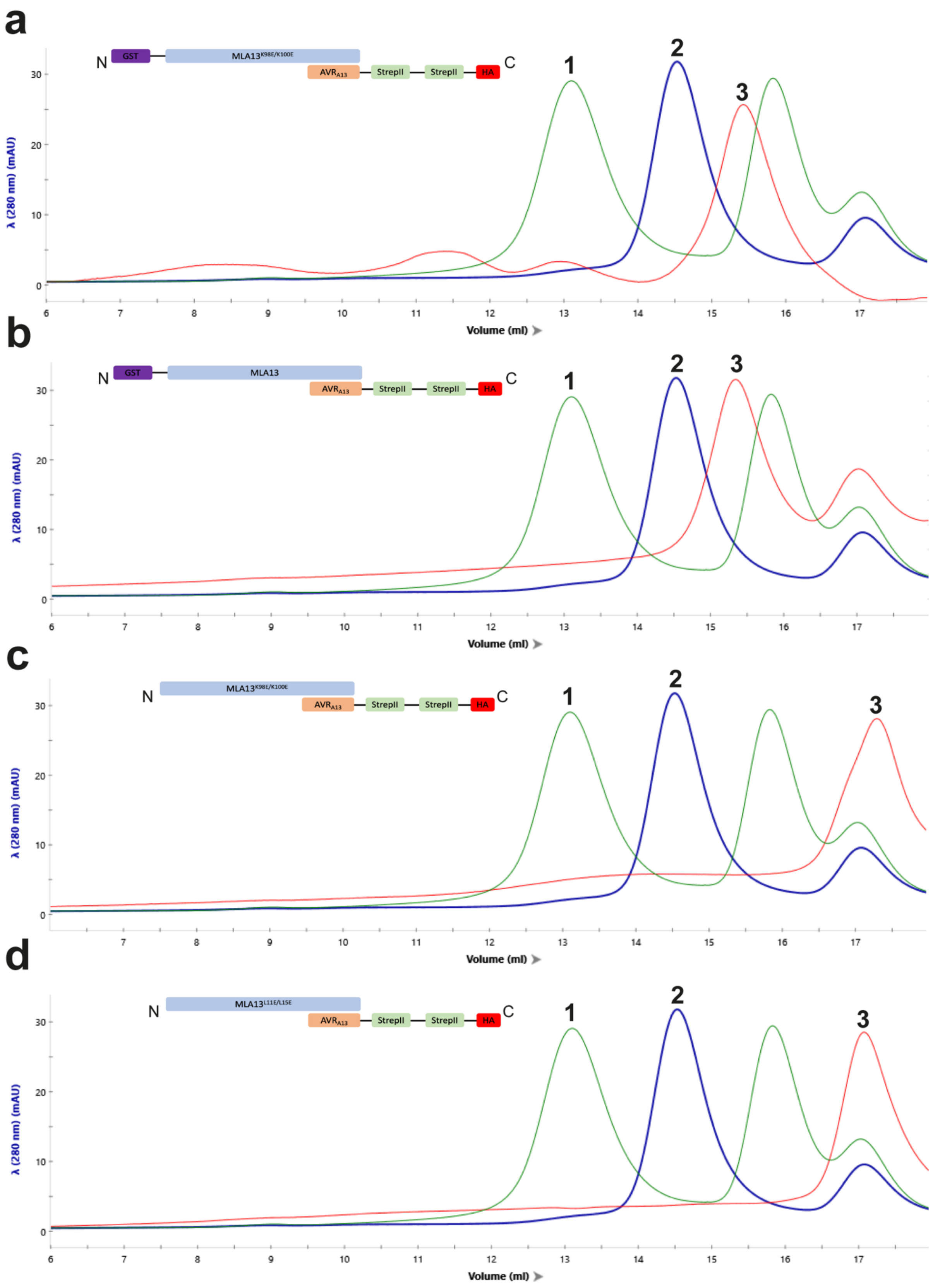


**Extended Data Fig. 7 | SEC profiles from purification experiments of various MLA13-$AVR_{A13}$-1 constructs.** SEC profiles of affinity-purified Sr35 (Sr35$^{L11E/L15E}$ (no tag) + AvrSr35-2Strep-HA; green trace) and Sr50 (Sr50$^{L11E/L15E}$ (no tag) + HA-2Strep-AvrSr50; blue trace) resistosomes. The blue and green resistosome traces are intended as size references for the traces of various MLA13-$AVR_{A13}$-1 constructs. Single-step affinity purification *via* the twin Strep tagged-effector followed by direct loading on SEC was used for all samples displayed. The Sr35 resistosome, Sr50 resistosome and MLA13-$AVR_{A13}$-1 heterocomplex are labelled as 1, 2 and 3, respectively. The MLA13-$AVR_{A13}$-1 heterocomplex consistently elutes at a later elution volume, indicating the extraction of a lower-order complex even under the exact same condition, tags and substitutions as Sr35 and Sr50 (see panel **d**).

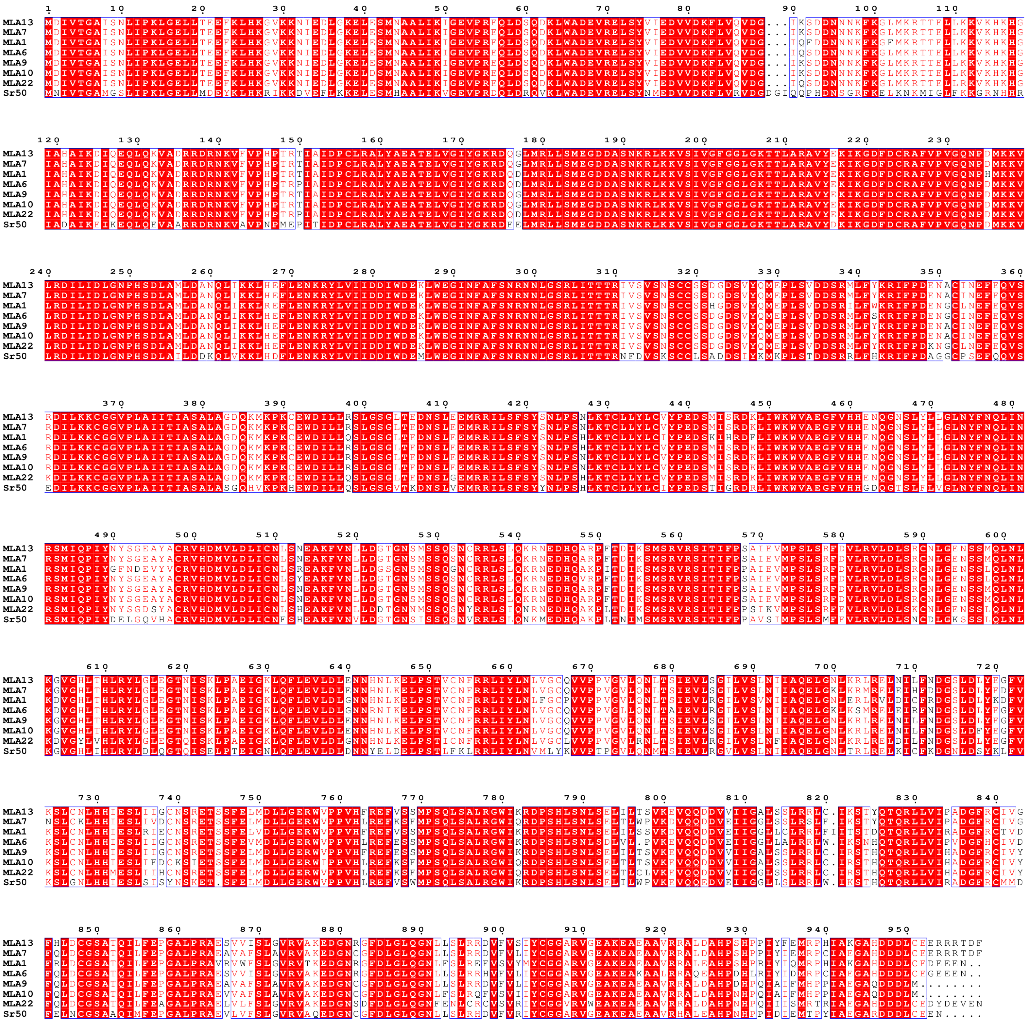

**Extended Data Fig. 8 | Amino acid sequence alignment of all the MLAs that recognise a matching *Bh* effector and Sr50.** Alignment performed using MUSCLE and visualised using ESPript 3.0.

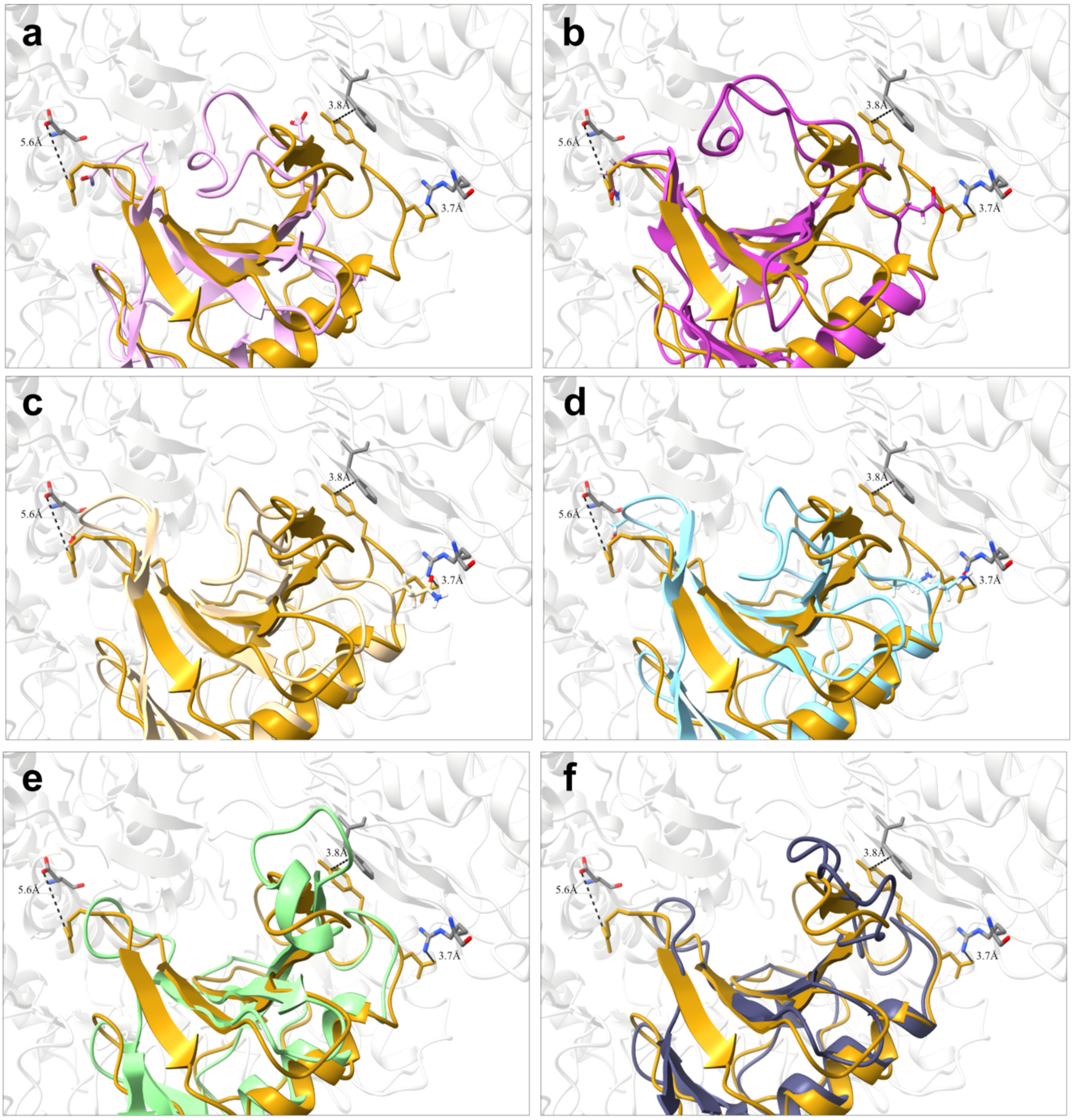


**Extended Data Fig. 9 | Structural alignment of $AVR_{A13}$-1 in the heterodimer with the crystal structures of other effectors.** Experimentally tested residues on $AVR_{A13}$-1 (dark goldenrod colour) that interact with MLA13 (transparent grey) are highlighted with proximity labels. Aligned effectors include **a,** $AVR_{A6}$, **b,** $AVR_{A7}$-2, **c,** $AVR_{A10}$, **d,** $AVR_{A22}$, **e,** CSEP0064, **f,** AvrPm2.

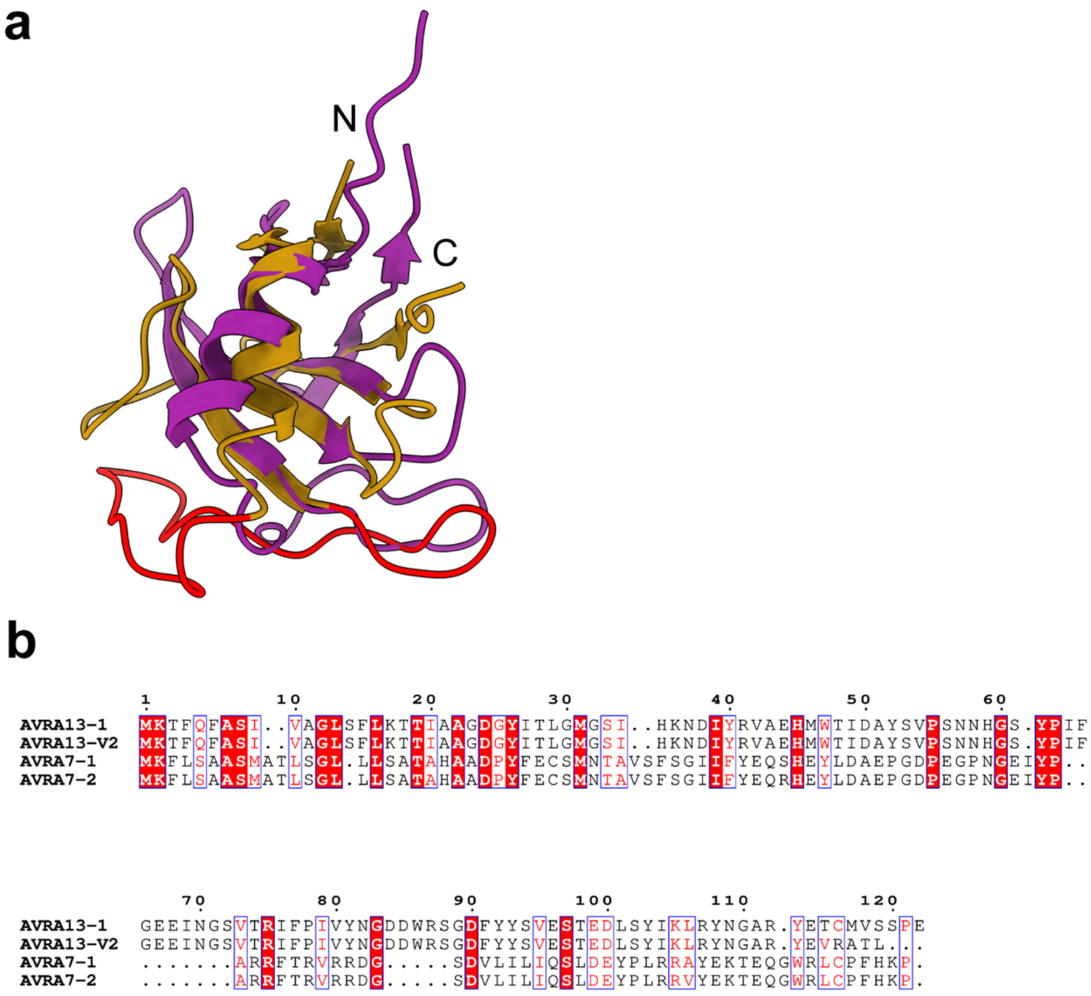


**Extended Data Fig. 10 | Structural and sequence alignments of AVR$_{A13}$ and AVR$_{A7}$ variants. a,** Structural alignment of AVR$_{A13}$-1 (dark goldenrod colour) and crystal structure of AVR$_{A7}$-1 (burgundy colour; PDB: 8OXL). The basal loops of AVR$_{A13}$-1 are coloured in red. **b,** Sequence alignment of AVR$_{A13}$ and AVR$_{A7}$ variants. Alignment performed using MUSCLE and visualised using ESPript 3.0.

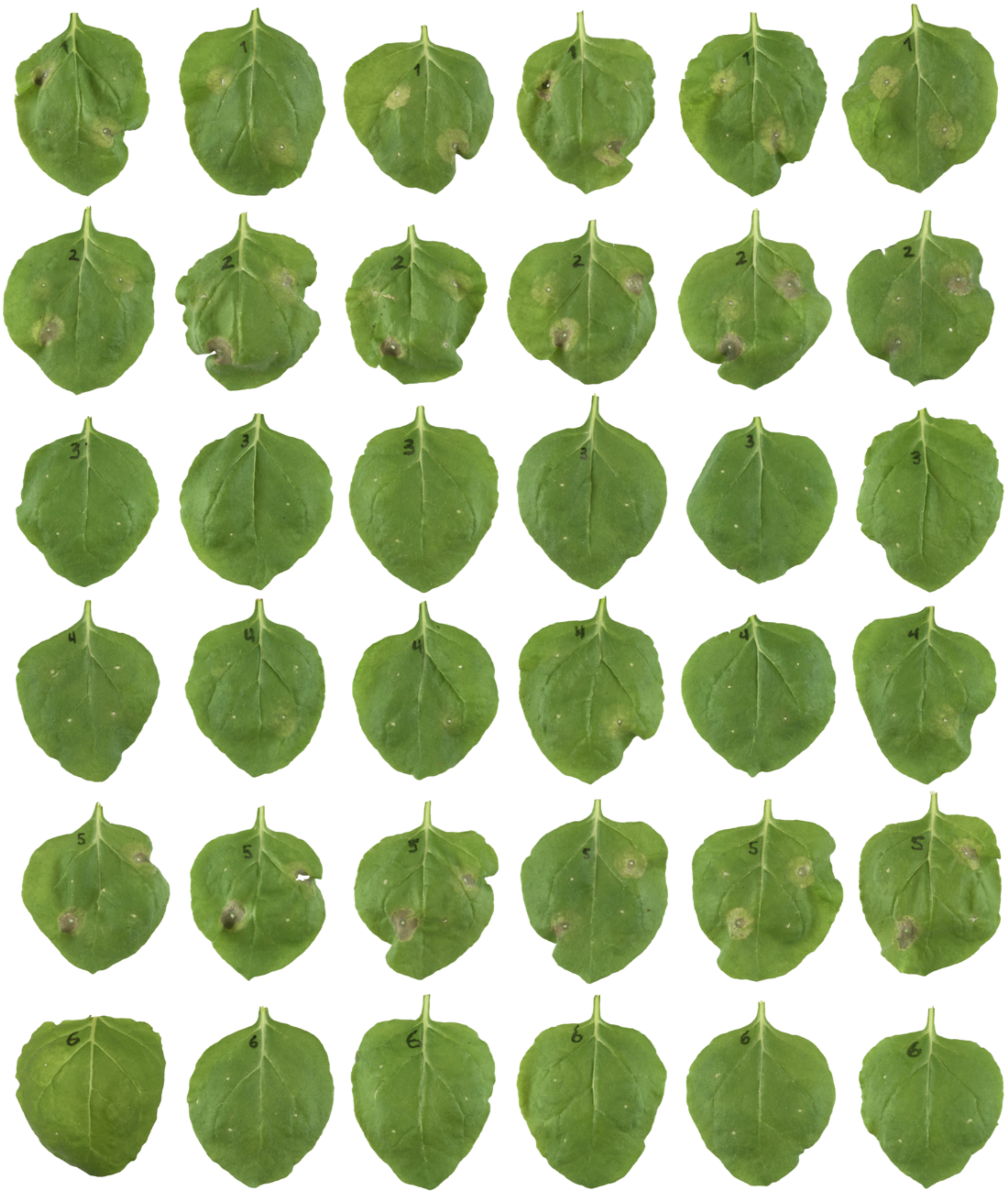

**Supplementary Fig. 1 | Six replicates of *Agrobacterium*-mediated co-expression of MLA13 variants with AVR$_{A13}$-1 and AVR$_{A22}$ in leaves of *N. benthamiana* from Extended Data Fig. 3.** Infiltration points described below are ordered sequentially counterclockwise starting in the upper left corner of each leaf series. Leaf #1: MLA13 + AVR$_{A13}$-1, MLA13$^{L11E/L15E}$ + AVR$_{A13}$-1, MLA13$^{F99E}$ + AVR$_{A13}$-1, MLA13$^{K98E/K100E}$ + AVR$_{A13}$-1; leaf #2: MLA13$^{K98E/F99E/K100E}$ + AVR$_{A13}$-1, MLA13$^{D502V}$ + AVR$_{A13}$-1, MLA13$^{L11E/L15E/D502V}$ + AVR$_{A13}$-1, MLA13$^{K98E/K100E/D502V}$ + AVR$_{A13}$-1; leaf #3: MLA13$^{K207R}$ + AVR$_{A13}$-1, GST- MLA13 + AVR$_{A13}$-1, GST- MLA13$^{D502V}$ + AVR$_{A13}$-1. Leaves 4-6 are the same as leaves 1-3 except co-expression of AVR$_{A22}$ replaces AVR$_{A13}$-1.

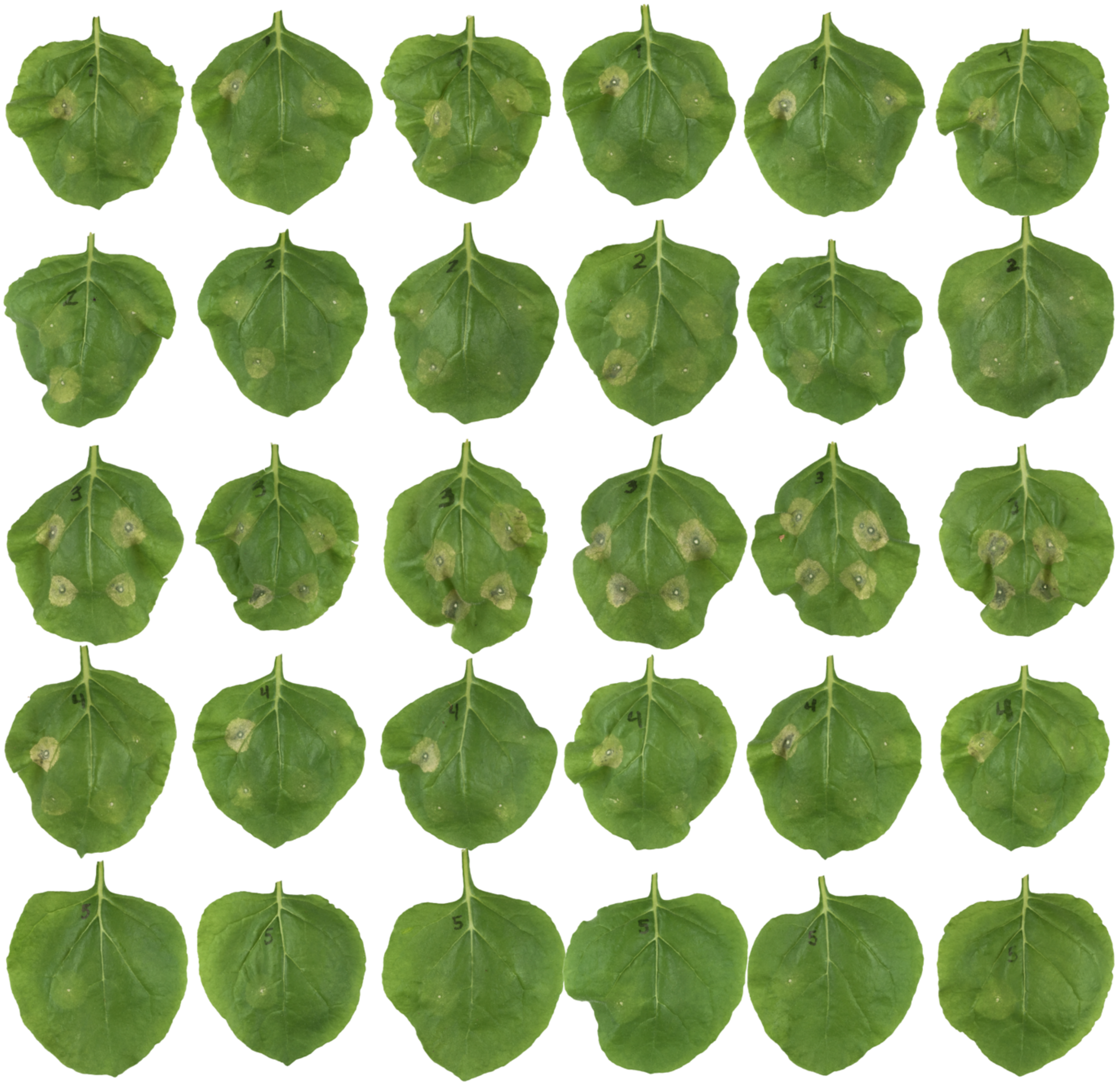

**Supplementary Fig. 2 | Six replicates of *Agrobacterium*-mediated co-expression of AVR$_{A13}$-1 variants with MLA13 in leaves of *N. benthamiana* from Fig.3c.** Infiltration points described below are ordered sequentially counterclockwise starting in the upper left corner of each leaf series. Leaf #1: AVR$_{A13}$-1 WT, AVR$_{A22}$ WT, Y52A, P55A; leaf #2: N58A, H59A, G60A, F65A; leaf #3: G66A, Y81A, N82A, Y108A; leaf #4: N109A, Y52A/G60A, Y52A/F65A, G60A/F65A; leaf #5: Y52A/G60A/F65A.

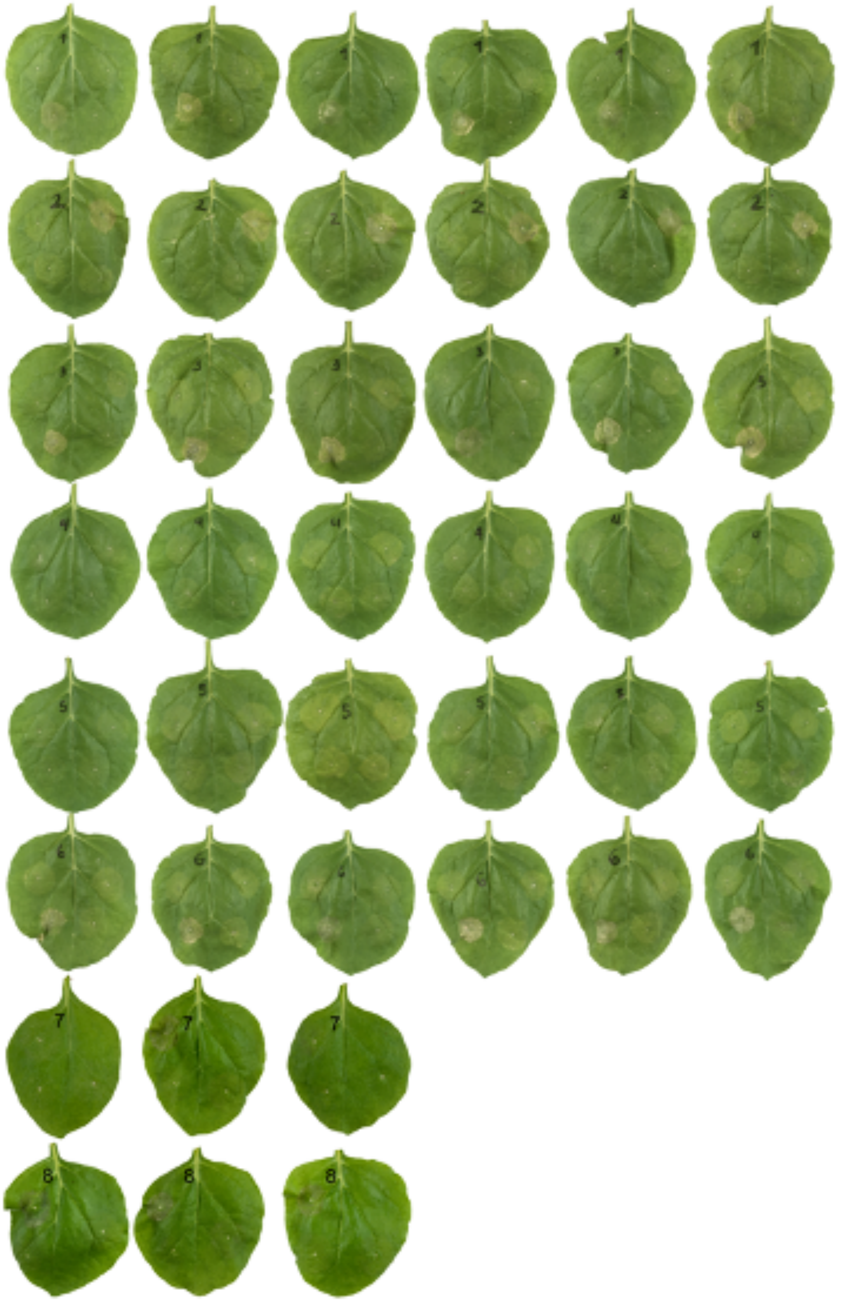

**Supplementary Fig. 3 | Replicates of *Agrobacterium*-mediated co-expression of MLA13 variants with AVR$_{A13}$-1 and AVR$_{A22}$ in leaves of *N. benthamiana* from Fig.4c.** Infiltration points described below are ordered sequentially counterclockwise starting in the upper left corner of each leaf series. Leaf #1: MLA13 WT + AVR$_{A22}$, MLA13 WT + AVR$_{A13}$-1, Y491A + AVR$_{A22}$, Y491A + AVR$_{A13}$-1; leaf #2: Y496A + AVR$_{A22}$, Y496A + AVR$_{A13}$-1, H643A +, AVR$_{A22}$, H643A + AVR$_{A13}$-1; leaf #3: S902A + AVR$_{A22}$, S902A + AVR$_{A13}$-1, Y934A +, AVR$_{A22}$, Y934A + AVR$_{A13}$-1; leaf #4: E936A + AVR$_{A22}$, E936A + AVR$_{A13}$-1, Y491A/H643A +, AVR$_{A22}$, Y491A/H643A + AVR$_{A13}$-1; leaf #5: H643A/E936A + AVR$_{A22}$, H643A/E936A + AVR$_{A13}$-1, Y491A/E936A +, AVR$_{A22}$, Y491A/E936A + AVR$_{A13}$-1; leaf #6: S902A/F935A + AVR$_{A22}$, S902A/F935A + AVR$_{A13}$-1, Y491A/H643A/E936A +, AVR$_{A22}$, Y491A/H643A/E936A + AVR$_{A13}$-1; relevant infiltration points on leaves 7 and 8 start at position two; leaf #7: F900A + AVR$_{A22}$, F900A + + AVR$_{A13}$-1; leaf #8: R938A + AVR$_{A22}$, R938A + AVR$_{A13}$-1.

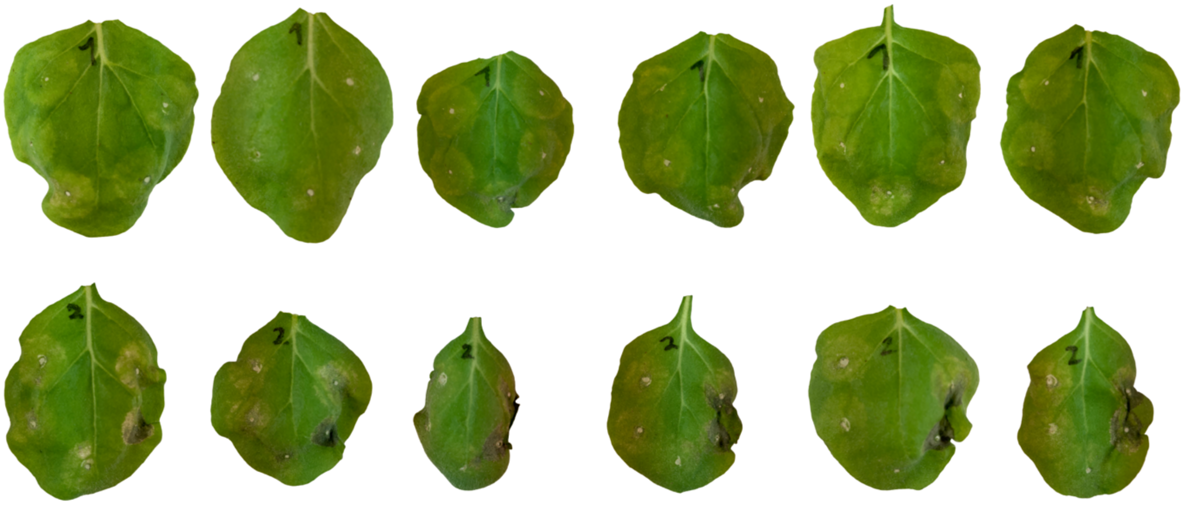

**Supplementary Fig. 4 | Replicates of *Agrobacterium*-mediated co-expression of MLA7 and MLA7$^{L902S}$ with effector variants in *N. benthamiana* as tested in Fig.5b.** Leaf #1 and leaf #2 are co-expressed with *Mla7* WT and *Mla7*$^{L902S}$, respectively. Infiltration points described below are ordered sequentially counterclockwise starting in the upper left corner of each leaf series: $AVR_{a22}$, $AVR_{a7}$-1, $AVR_{a7}$-2, $AVR_{a13}$-1, $AVR_{a13}$-V2.

**Supplementary Table 1 | Statistical output from cryo-EM data processing and structural model building.**

| | | |
|---|---|---|
| **Sample conditions** | | |
| Grid type | Quantifoil Cu R2/4 (200 mesh) + Graphene Oxide | |
| **Cryo-EM data collection** | | |
| Microscope | Titan Krios G3i | |
| Voltage (kV) | 300 | |
| Spherical aberration (mm) | 2.7 | |
| Condenser C2 aperture (µm) | 70 | |
| Objective aperture size (µm) | 100 | |
| Camera | Falcon III | |
| Pixel size | 0.862 | |
| Total dose (electron*Å$^{-2}$) | 42 | |
| Number of frames | 42 | |
| Images per hole | 3 | |
| Energy filter | None | |
| Defocus range (µm) | -2.0 to -0.3 | |
| # micrographs collected | 8,188 | |
| # micrographs used | 5,897 | |
| **Cryo-EM data processing** | | |
| software | cryosparc v4.4.1+patch240110 | |
| Particles | | |
| after 2D classification | 115,686 | |
| after 3D sorting | 48,191 | |
| Resolution (FSC 0.143, Å) | 3.8 | |
| **Model building and refinement** | | |
| Software for building | Coot 0.9.4.7 EL | |
| Residues build | | |
| MLA13 | 2-131,143-541,555-956 | |
| AVRa13 | 25-122 | |
| Software for refinement | PHENIX 1.21 - 5207 | |
| Composition (#) | | |
| Chains | 2 | |
| Atoms | 8127 (Hydrogens: 0) | |
| Residues | Protein: 1029 | |
| Water | 0 | |
| Ligands | 0 | |
| Bonds (RMSD) | | |
| Length (Å) (# > 4σ) | 0.002 (0) | |
| Angles (°) (# > 4σ) | 0.518 (1) | |
| MolProbity score | 2.04 | |
| Clash score | 8.96 | |
| Ramachandran plot (%) | | |
| Outliers | 0.10 | |
| Allowed | 10.19 | |
| Favored | 89.72 | |
| Rama-Z (Z-score, RMSD) | | |
| whole (N = 1021) | -2.06 (0.26) | |
| helix (N = 331) | -0.17 (0.29) | |
| sheet (N = 156) | -0.63 (0.46) | |
| loop (N = 534) | -2.39 (0.25) | |
| Rotamer Outliers (%) | 0.0 | |
| Peptide plane (%) | | |
| Cis proline/general | 2.8/0.0 | |
| Twisted proline/general | 0.0/0.0 | |
| CaBLAM outliers (%) | 6.81 | |
| Supplied Resolution (Å) | 3.5 | |
| Resolution Estimates (Å) | Masked | Unmasked |
| d 99 (full) | 2.5 | 2.5 |
| d model | 2.2 | 2.2 |
| d FSC model (0/0.143/0.5) | 1.7/2.0/3.7 | 1.7/2.0/3.7 |
| Model vs. Data | | |
| CC (mask | 0.71 | |
| CC (box) | 0.71 | |
| CC (peaks) | 0.69 | |
| CC (volume) | 0.72 | |
